# Transcriptomic profiling reveals distinct subsets of immune checkpoint inhibitor-induced myositis

**DOI:** 10.1101/2022.12.12.520136

**Authors:** Iago Pinal-Fernandez, Angela Quintana, Jose C. Milisenda, Maria Casal-Dominguez, Sandra Muñoz-Braceras, Assia Derfoul, Jiram Torres-Ruiz, Katherine Pak, Stefania Del Orso, Faiza Naz, Gustavo Gutierrez-Cruz, Margherita Milone, Shahar Shelly, Yaiza Duque-Jaimez, Ester Tobias-Baraja, Ana Matas-Garcia, Gloria Garrabou, Joan Padrosa, Javier Ros, Ernesto Trallero-Araguás, Brian Walitt, Lisa Christopher-Stine, Thomas E. Lloyd, Chen Zhao, Shannon Swift, Arun Rajan, Josep Maria Grau, Albert Selva-O’Callaghan, Teerin Liewluck, Andrew L. Mammen

**Affiliations:** Muscle Disease Unit, National Institute of Arthritis and Musculoskeletal and Skin Diseases, National Institutes of Health, Bethesda, MD, USA; Department of Neurology, Johns Hopkins University School of Medicine, Baltimore, MD, USA; Systemic Autoimmune Disease Unit, Vall d’Hebrón Institute of Research, Barcelona, Spain; Muscle Research Unit, Internal Medicine Service, Hospital Clinic, Barcelona, Spain; CIBERER, Barcelona, Spain; Division of Neuromuscular Medicine, Department of Neurology, Mayo Clinic, Rochester, MN, USA; Department of Neurology, Rambam Health Care Campus, Bruce Rappaport Faculty of Medicine, Technion-Israel Institute of Technology, Haifa, Israel; Medical Oncology, Vall d’Hebrón Hospital, Barcelona, Spain; Rheumatology Department, Vall d’Hebron Hospital, Barcelona, Spain; Division of Intramural Research, Department of Health and Human Services, National Institute of Nursing Research, National Institutes of Health, Bethesda, MD, USA; Department of Medicine, Division of Rheumatology, Johns Hopkins University School of Medicine, Baltimore, MD, USA; Thoracic and Gastrointestinal Malignancies Branch, Center for Cancer Research, National Cancer Institute, National Institutes of Health, Bethesda, MD, USA; Universitat Autónoma de Barcelona, Barcelona, Spain

**Author notes:** These authors contributed equally to this project (Iago Pinal-Fernandez, Angela Quintana, Jose C. Milisenda, Maria Casal-Dominguez). These authors contributed equally to this project (Teerin Liewluck, Andrew L. Mammen). **Address correspondence to**: Andrew L. Mammen, M.D., Ph.D., or Iago Pinal-Fernandez, M.D., Ph.D. Muscle Disease Unit, Laboratory of Muscle Stem Cells and Gene Regulation, National Institute of Arthritis and Musculoskeletal and Skin Diseases, National Institutes of Health, 50 South Drive, Room 1141, Building 50, MSC 8024, Bethesda, MD 20892. or.

**Keywords:** Immune checkpoint inhibitors, myositis, immune-related adverse events, anti-PD-1, anti-PD-L1, IL6, IFNG, interferon

## Abstract

**Objectives:** Inflammatory myopathy or myositis is a heterogeneous family of immune-mediated diseases including dermatomyositis (DM), antisynthetase syndrome (AS), immune-mediated necrotizing myopathy (IMNM), and inclusion body myositis (IBM). Immune checkpoint inhibitors (ICI) can also cause myositis (ICI-myositis). This study was designed to define gene expression patterns in muscle biopsies from patients with ICI-myositis.

**Methods:** Bulk RNA sequencing was performed on 200 muscle biopsies (35 ICI-myositis, 44 DM, 18 AS, 54 IMNM, 16 IBM, and 33 normal muscle biopsies) and single nuclei RNA sequencing was performed on 22 muscle biopsies (7 ICI-myositis, 4 DM, 3 AS, 6 IMNM, and 2 IBM).

**Results:** Unsupervised clustering defined three distinct transcriptomic subsets of ICI-myositis: ICI-DM, ICI-MYO1, and ICI-MYO2. ICI-DM included patients with DM and anti-TIF1γ autoantibodies who, like DM patients, overexpressed type 1 interferon-inducible genes. ICI-MYO1 patients had highly inflammatory muscle biopsies and included all patients that developed co-existing myocarditis. ICI-MYO2 was composed of patients with predominant necrotizing pathology and low levels of muscle inflammation. The type 2 interferon pathway was activated both in ICI-DM and ICI-MYO1. Unlike the other types of myositis, all three subsets of ICI-myositis patients overexpressed genes involved in the IL6 pathway.

**Conclusions:** We identified three distinct types of ICI-myositis based on transcriptomic analyses. The IL6 pathway was overexpressed in all groups, the type I interferon pathway activation was specific for ICI-DM, the type 2 IFN pathway was overexpressed in both ICIDM and ICI-MYO1, and only ICI-MYO1 patients developed myocarditis.

## Introduction

Inflammatory myopathy or myositis is a heterogeneous family of immune-mediated diseases that includes dermatomyositis (DM), antisynthetase syndrome (AS), immune-mediated necrotizing myopathy (IMNM), and inclusion body myositis (IBM).[1] Each of these is associated with distinctive clinical features and muscle biopsies from each type of myositis have unique histopathological features and gene expression profiles.[2]

In recent years, it has been recognized that immune checkpoint inhibitors (ICI) can trigger a novel form of myositis in cancer patients (ICI-myositis). Muscle biopsies from ICI-myositis patients are characterized by the presence of T cells and macrophages with minimal B cell infiltration.[3–6] Unlike patients with DM, AS, IMNM, or IBM, ICI-myositis is often accompanied by myasthenia gravis or a myasthenia gravis-like phenotype,[7] and/or myocarditis. Furthermore, a minority of patients with ICI myositis also have a DM-like skin rash. However, it remains unknown whether muscle biopsies from ICI myositis patients have a unique transcriptomic profile and whether biopsies from those patients with and without DM-like rashes have different gene expression patterns. In this study, we addressed these questions by performing bulk and single-nuclei RNA sequencing on muscle biopsy tissue obtained from patients with ICI-myositis and comparing them to other types of myositis.

## Methods

### Patients and samples

Thirty-five patients from the National Institutes of Health (n=4, Bethesda, US), Mayo Clinic (n=11, Rochester, US), Vall d’Hebron Hospital (Barcelona, Spain), and Clinic Hospital (n=20, Barcelona, Spain) with a myopathy related to ICI treatment (either anti-PD-1 or anti-PD-L1 therapy alone or in combination with anti-CTLA-4) and an available frozen muscle biopsy were included in the study. ICI-myositis was defined as the presence of muscle weakness, or the presence of myopathic changes with or without prominent inflammatory infiltrate in the muscle biopsy after starting treatment with ICI.

The epidemiologic features of the patients, type and cycle of ICI, clinical features (including the presence or absence of myocarditis, dysphagia, and ocular involvement), autoantibody profile (including myositis-specific [anti-HMGCR, anti-SRP, anti-Mi2, anti-NXP2, anti-TIF1g, anti-MDA5, anti-Jo1], myositis-associated autoantibodies [anti-Ro52, anti-NT5c1a, anti-PM/Scl], anti-AChR, anti-Musk, and anti-striational autoantibodies) were included in the analysis.

The biopsies from patients with ICI-myositis were compared to 33 normal muscle biopsies and 132 muscle biopsies from patients with the most common types of inflammatory myopathy, including 54 IMNM, 44 DM, 18 AS, and 16 IBM. Of note, the transcriptomic profiles of the DM, AS, IMNM, and IBM muscle biopsies have been previously reported.[2, 8–11] All biopsies were from subjects enrolled in institutional review board (IRB)-approved longitudinal cohorts in the different hospitals.

The transcriptional characteristics of the muscle biopsies were also compared with publicly available transcriptomic datasets of tumors before and after ICI treatment (GEO: GSE91061).[12]

### Differentiating human skeletal muscle myoblasts treated with different types of interferon

We treated human skeletal muscle myoblasts (HSMMs) to identify the IFNB1 and IFNG-specific interferon-stimulated genes. Normal HSMMs were cultured according to recommended protocol by the manufacturer (Lonza). When 80% confluent, the cultures were induced to differentiate into myotubes by replacing the growth medium with differentiation medium (Dulbecco’s modified Eagle’s medium supplemented with 2% horse serum and L-glutamine). Differentiating HSMMs were treated daily with 100U/L and 1000U/L of IFNA2a (R&D, ref:11100-1), IFNB1 (PeproTech, ref:300-02BC), or IFNG (PeproTech, ref:300-02) for 7 days and then harvested for RNA extraction and subsequent RNA sequencing.

### Bulk RNA sequencing

Bulk RNA sequencing was performed on frozen muscle biopsy specimens as previously described.[2, 8–10] In short, RNA was extracted from fresh-frozen biopsies using TRIzol (Invitrogen) and quantified using NanoDrop. Libraries for bulk RNA sequencing were prepared using NEBNext Poly(A) mRNA Magnetic Isolation Module and Ultra^™^ II Directional RNA Library Prep Kit for Illumina (New England BioLabs, cat. #E7490 and #E7760). The input RNA and the resulting libraries were analyzed with Agilent 4200 Tapestation for quality assessment. The libraries were sequenced using the NextSeq 550 and the NovaSeq 6000 Illumina platforms.

### Single-nuclei RNA-sequencing

We performed single-nuclei RNA sequencing in 4 ICI-myositis, 3 ICI-dermatomyositis (ICIDM), 4 DM, 3 AS, 6 IMNM, and 2 IBM. For the nuclei isolation, we used a modification of the sucrose-gradient ultracentrifugation nuclei isolation protocol from Schirmer et al.[13] Ten mg of frozen muscle tissue was sectioned and homogenized in 1mL of lysis buffer (0.32M sucrose, 5mM CaCl2, 3mM MgCl2, 0.1mM EDTA, 10mM Tris-HCl pH 8, 1mM DTT, 0.5% Triton X-100 in DEPC-treated water) using 1.4mm ceramic beads low-binding tubes and the Bertin Technology Precellys 24 lysis homogenizer (6500rpm-3times x 30s). The homogenized tissue was transferred into open-top thick-walled polycarbonate ultracentrifuge tubes (25×89 mm, Beckman Coulter) on ice. 3.7mL of sucrose solution (1.8 M sucrose, 3mM MgCl2, 1mM DTT, 10mM Tris-HCl) were pipetted to the bottom of the tube containing lysis buffer generating two separated phases (sucrose on the bottom and homogenate on the top). The tubes were filled almost completely with lysis buffer and weighted for balance. The samples were ultracentrifuged (Beckman Coulter XE-90, SW32 rotor, swinging bucket) at 24,400rpm (107,163rcf) for 2.5 hours at 4°C, transferred to ice, and the supernatant removed. Two hundred microliters of DEPC-PBS were added to each pellet, incubated on ice for 20 minutes, and then pellets were resuspended. The resulting samples were filtered twice using 30*μ*m Miltenyi pre-separation filters. The nuclei were counted using a manual hemocytometer. Between 2000 and 3000 nuclei per sample were loaded in the 10X Genomic Single-Cell 3’ system. We performed the 10X nuclei capture and the library preparation protocol according to the manufacturer’s instructions without modification.

### Statistical analysis

Bulk RNAseq reads were demultiplexed using bcl2fastq/2.20.0 and preprocessed using fastp/0.21.0. The abundance of each gene was generated using Salmon/1.5.2 and quality control output was summarized using multiqc/1.11. Dimensionality reduction was performed with the uniform manifold approximation and projection (UMAP) using umap/0.2.9.0. The number of clusters was determined with Tibshirani’s gap statistic[14] using factoextra/1.0.7. The clusters were defined using the K-means algorithm. Counts were normalized using the Trimmed Means of M values (TMM) from edgeR/3.34.1 for graphical analysis. Differential expression was performed using limma/3.48.3. Pathway analysis was done using Gene Set Enrichment Analysis (GSEA) using clusterProfiler/4.6.0 and GSEA/4.2.3 for the Reactome and the Hallmark datasets.[15]

For the single-cell and single-nuclei RNAseq, reads were demultiplexed and aligned using cellranger/6.0.1. The samples were aggregated, normalized (SCTransform), and integrated (RunHarmony) using Seurat/4.1.0. Graphical analysis of single cell and single nuclei RNAseq data used the functions contained in Seurat/4.1.0.

The Benjamini-Hochberg correction was used to adjust for multiple comparisons and a corrected p-value (q-value) of 0.05 or less was considered statistically significant. Graphical analysis used both the Python and R programming languages.

## Results

### Clinical features of patients with ICI-myositis

Thirty-five patients (11 female, average age 67yo) with ICI-myositis were included in the study and three (8.6%) of these had DM rashes. Twenty-four (69%) were treated with PD1 inhibitors, and 11 (31%) with PD-L1 inhibitors. In addition, five (14%) were concomitantly treated with CTLA-4 inhibitors. The three most prevalent primary tumors were melanoma (n=10), thymoma (n=5), and lung cancer (n=4). Sixteen (46%) developed myopathy after receiving their first cycle of ICI, and the rest of them after two or more cycles. Seven ICI-myopathy patients (20%) had diplopia and 13 (37%) of them developed myocarditis. Twenty-nine (83%) patients had autoantibodies against the neuromuscular junction or skeletal muscle antigens: 14 had anti-acetylcholine receptor (AChR) antibodies, 15 had anti-striational antibodies, and 16 were positive for various other myositis autoantibodies. The three patients with DM rashes had high titers of anti-TIF1γ autoantibodies. In three patients with pre-ICI treatment serum available, the same autoantibodies that were present after treatment with ICI were detectable before ICI initiation (two patients with anti-AChR and anti-striational autoantibodies and one patient with DM rashes and anti-TIF1*γ* autoantibodies). The complete characteristics of these patients and their clinical evolution are described in Table 1 and Supplemental Table S1. The clinical features of the DM, AS, IMNM, and IBM subjects have been previously reported.[2, 8–11]

**Table 1.** General features of immune-checkpoint induced myopathy groups resulting from unsupervised clustering (ICI-DM, ICI-MYO1, ICI-MYO2).

| Characteristic | Overall, N = 35<sup>1</sup> | ICI-DM, N = 3<sup>1</sup> | ICI-MYO1, N = 24<sup>1</sup> | ICI-MYO2, N = 8<sup>1</sup> | p-value<sup>2</sup> |
| --- | --- | --- | --- | --- | --- |
| Female | 11 (31%) | 1 (33%) | 8 (33%) | 2 (25%) | >0.9 |
| Age at biopsy | 67 (60, 74) | 73 (58, 74) | 64 (60, 72) | 69 (67, 72) | 0.6 |
| Dermatomyositis | 3 (8.6%) | 3 (100%) | 0 (0%) | 0 (0%) | **<0.001** |
| Inflammatory infiltrates | 25 (71%) | 3 (100%) | 19 (79%) | 3 (38%) | 0.056 |
| Treatment cycle |  |  |  |  | >0.9 |
| 1 | 16 (46%) | 1 (33%) | 11 (46%) | 4 (50%) |  |
| 2 | 13 (37%) | 2 (67%) | 8 (33%) | 3 (38%) |  |
| 3 | 5 (14%) | 0 (0%) | 4 (17%) | 1 (12%) |  |
| 4 | 1 (2.9%) | 0 (0%) | 1 (4.2%) | 0 (0%) |  |
| PD1 inhibitor | 24 (69%) | 2 (67%) | 14 (58%) | 8 (100%) | 0.065 |
| PD-L1 inhibitor | 11 (31%) | 1 (33%) | 10 (42%) | 0 (0%) | 0.065 |
| CTLA4 inhibitor | 5 (14%) | 0 (0%) | 2 (8.3%) | 3 (38%) | 0.11 |
| Myocarditis | 13 (37%) | 0 (0%) | 13 (54%) | 0 (0%) | **0.006** |
| Diplopia | 7 (20%) | 0 (0%) | 6 (25%) | 1 (12%) | 0.8 |
| Dysphagia | 4 (11%) | 0 (0%) | 2 (8.3%) | 2 (25%) | 0.5 |
| Anti-striational | 15 (43%) | 0 (0%) | 11 (46%) | 4 (50%) | 0.6 |
| Anti-AChR | 14 (40%) | 0 (0%) | 12 (50%) | 2 (25%) | **0.010** |
| Other myositis autoantibodies | 16 (46%) | 3 (100%) | 12 (50%) | 1 (12%) | 0.051 |
| CK at biopsy | 652 (219, 1,218) | 428 (278, 4,758) | 896 (278, 1,394) | 427 (62, 830) | 0.3 |
| Peak CK | 1,184 (462, 6,118) | 428 (278, 4,758) | 1,240 (899, 6,565) | 652 (379, 4,771) | 0.4 |

<sup>1</sup>n (%); Median (IQR)
<sup>2</sup>Fisher's exact test; Kruskal-Wallis rank sum test

### Unsupervised clustering reveals three transcriptomically distinct groups of ICI-myositis patients

To determine whether distinct subtypes of ICI-myositis could be defined based on transcriptomic data from muscle biopsies, we performed unsupervised clustering analysis using the expression levels of genes in the different muscle biopsies. This revealed three distinct clusters of biopsies and patients in each cluster had unique clinical features as well. The ICI-DM cluster (n=3) included the three patients with DM rashes (p<0.001). The ICI-MYO1 cluster (n=24) included all the patients who developed myocarditis and patients in this cluster had a higher prevalence of anti-AChR autoantibodies (50% vs. 25%) compared to patients in the ICI-MYO2 cluster (n=8) (Figure 1, Table 1, Supplementary Table 1).

**Figure 1.**
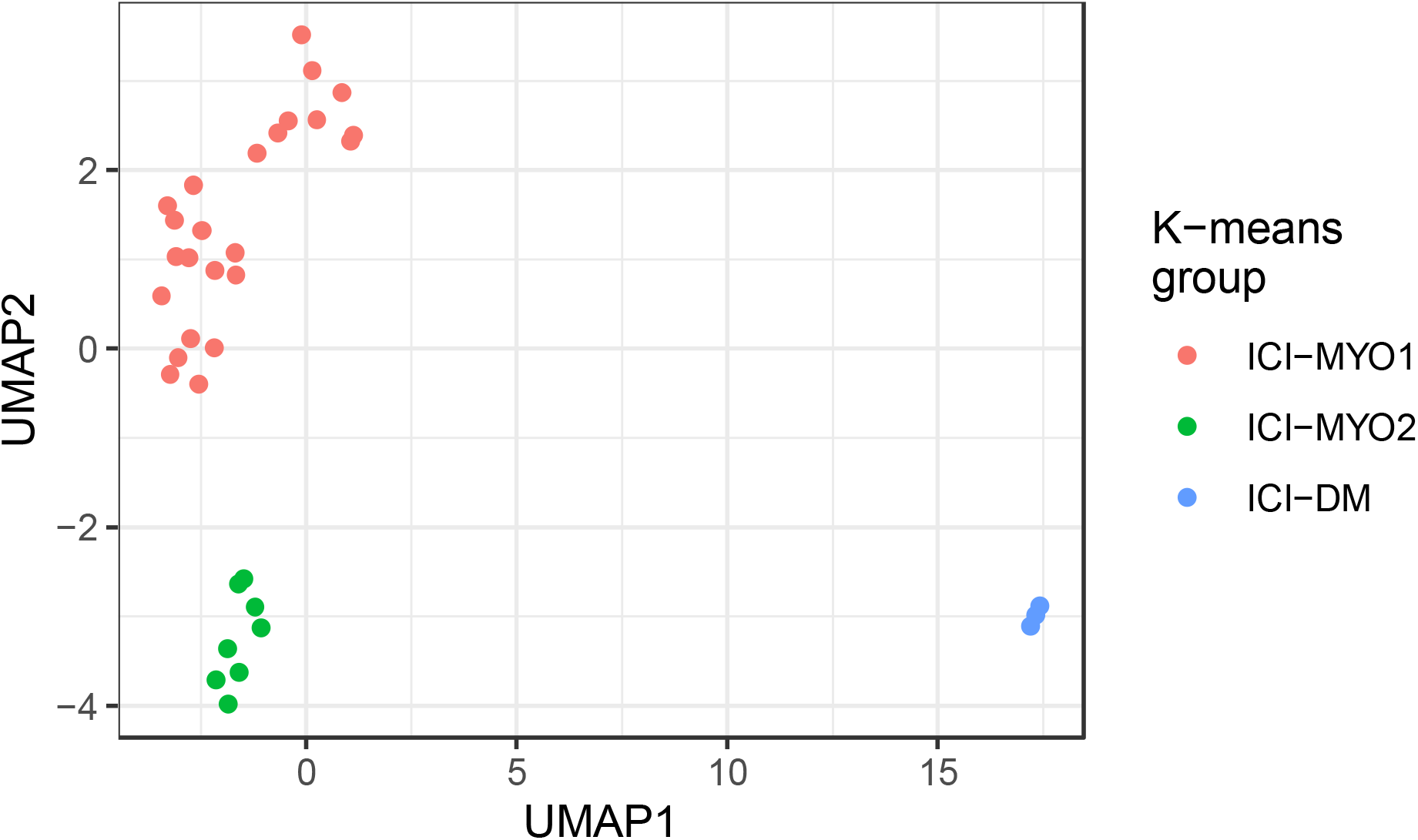
Groups of patients (ICI-MYO1, ICI-MYO2, ICI-DM) resulting from applying unsupervised clustering to the bulk RNA sequencing data of patients with immune checkpoint-induced (ICI) myopathy. Uniform manifold approximation and projection (UMAP) was used to perform the clustering, the number of clusters was determined using the gap statistic and the clusters were defined using the K-means algorithm.

Muscle biopsies from two patients in the ICI-DM cluster had perifascicular atrophy and the third had myofiber necrosis with intense perivascular inflammation. Muscle biopsies from patients in the ICI-MYO1 cluster were highly inflammatory compared to those in the ICI-MYO2 cluster, where necrosis was the predominant muscle biopsy feature. The inflammatory infiltrates of biopsies from the ICI-MYO1 cluster were predominantly composed of macrophages and CD8+ cells (Figure 2).

**Figure 2.**
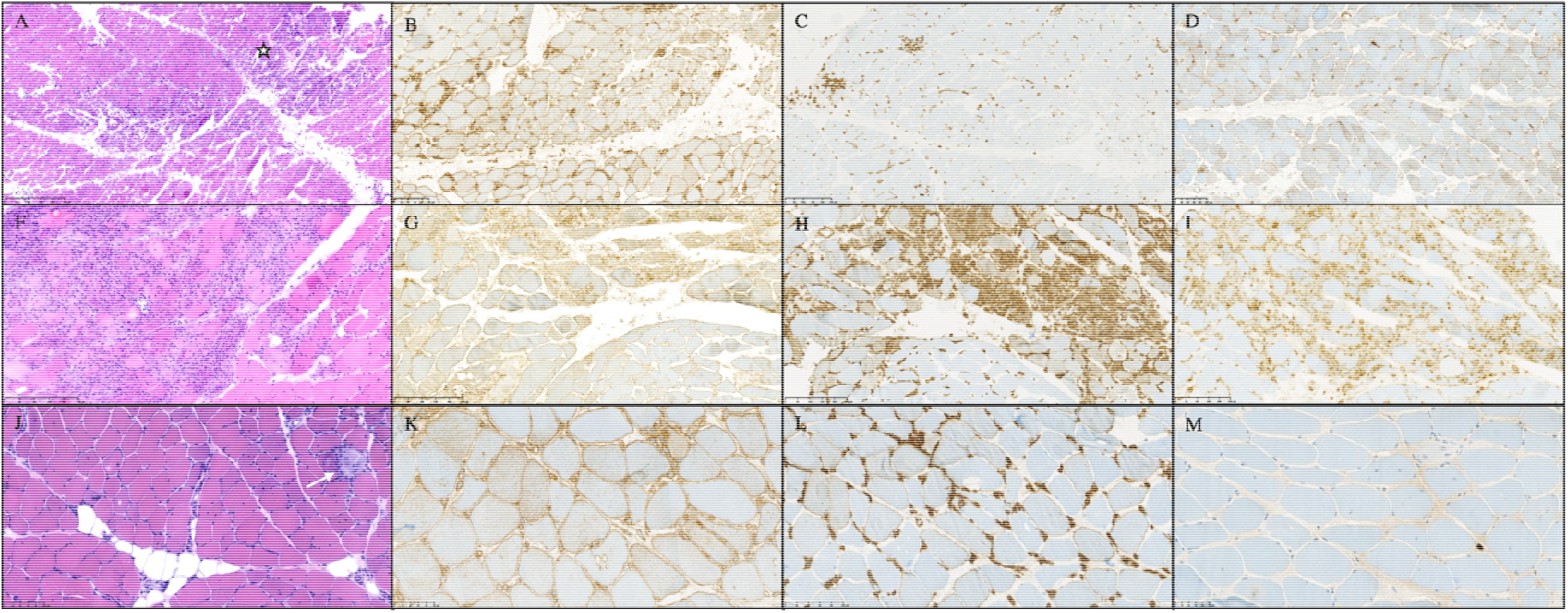
Histological appearance of patients with immune-checkpoint induced myopathy from clusters ICI-DM (A-D), ICI-MYO1 (F-I), and ICI-MYO2 (J-M). ICI-DM biopsies showed perifascicular atrophy, and intense vascular damage (star marks an area of muscle infarction). ICI-MYO1 showed intense inflammatory infiltrates with abundant macrophages and T-cells. Finall, ICI-MYO2 had predominant necrosis (white arrow indicates an area of myophagocytosis) with few inflammatory cells. The first column shows H&E (A, F, J), the second MHC-1 (B, G, K), the third MHC-2 (C, H, L). The fourth column shows CD68 in I and MX1 in D and M.

All the single-nuclei RNAseq analyses were performed using muscle biopsies from patients in the ICI-MYO1 cluster (Supplementary Figure 1).

### Type 2 interferon-inducible genes are overexpressed in ICI-MYO1 and ICI-DM

IFNγ and IFNγ-inducible genes were robustly overexpressed in patients with both ICI-MYO1 (e.g. GBP2 log2 fold-change[FC] 2.5, q-value 1.2e-13) and ICI-DM (e.g. GBP2 log2 fold-change[FC] 3, q-value 1.2e-11). The expression levels of these genes were comparable to that observed in patients with AS, and IBM. Biopsies in the ICI-MYO2 cluster were significantly lower than in ICI-DM or ICI-MYO1, but still had significantly higher levels of these genes compared to normal biopsies (GBP2 log2 fold-change[FC] 1.2, q-value 6e-6) (Figure 3–4, Supplementary Figure 2-4, Supplementary Table 3).

**Figure 3.**
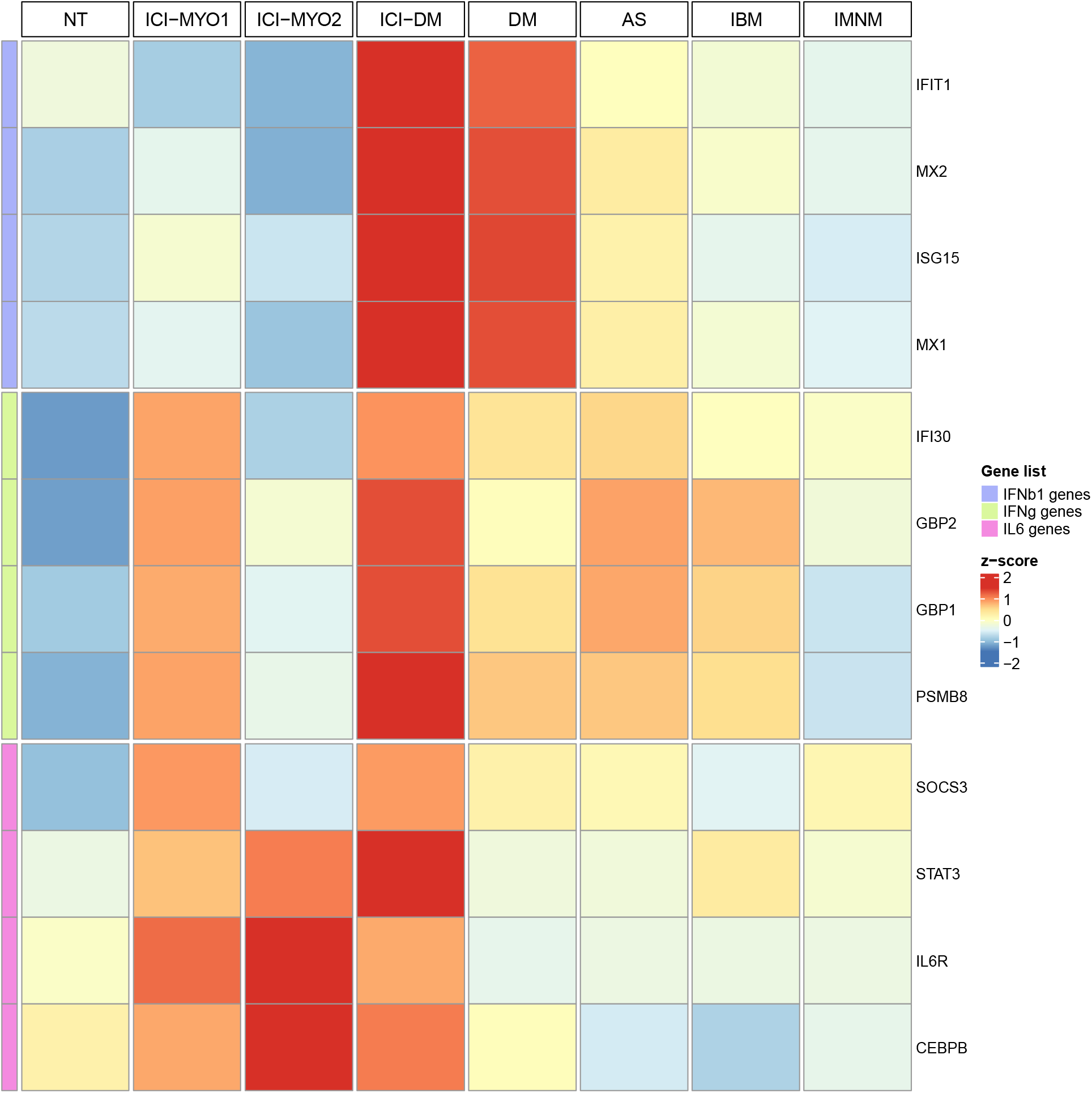
Expression (average z-score of log2[TMM+1]) of IFNB1, IFNG, and IL6 related genes in the three clusters of patients with ICI-induced myopathy (ICI-MYO1, ICI-MYO2, and ICI-DM) and in the comparator muscles biopsies. NT: normal muscle; DM: dermatomyositis; AS: antisynthetase syndrome; IBM: inclusion body myositis; IMNM: immune-mediated necrotizing myopathy. These IFNB1 and IFNG inducible genes were validated in cultures of differentiating human skeletal muscle myoblasts treated with IFNB1 and IFNG.

**Figure 4.**
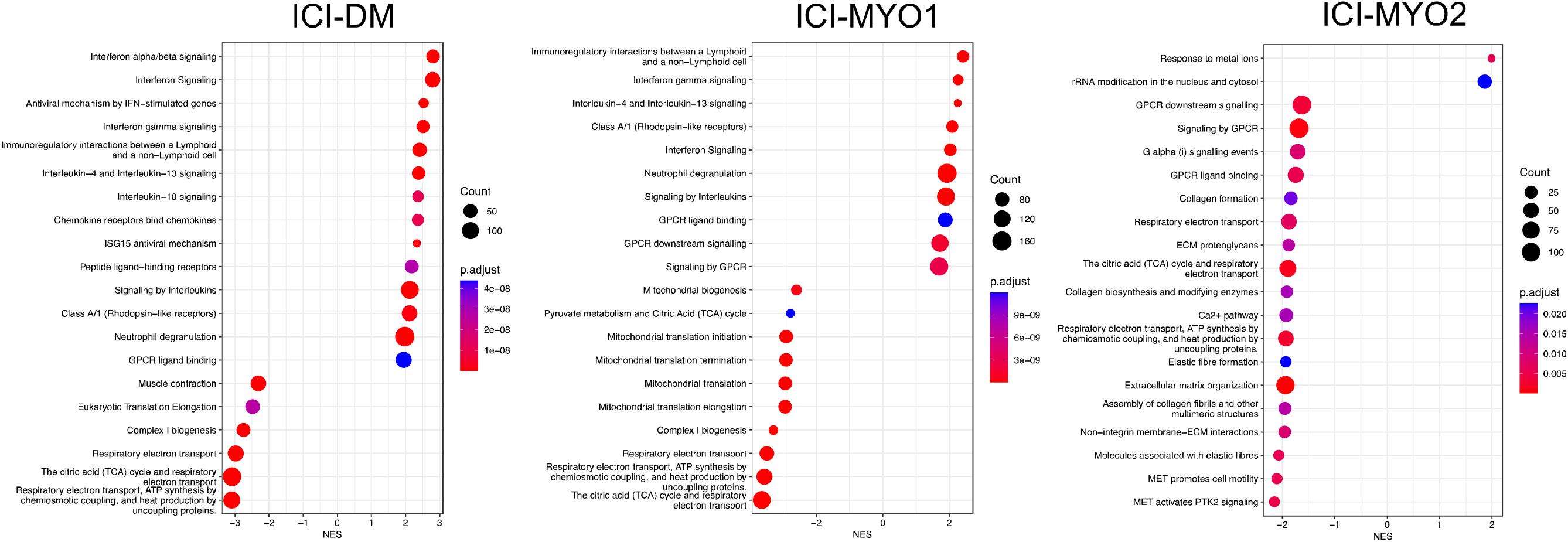
Top pathways in gene set enrichment analysis using the Reactome database in the three clusters of patients with ICI-induced myopathy (ICI-DM [left], ICI-MYO1 [middle], and ICI-MYO2 [right]).

Although single-nuclei RNA-seq identified minimal levels of IFNG in T-cells, genes specifically upregulated by IFNG were, in general, below the detection threshold of this technique (Figure 5, Supplementary Table 4-5).

**Figure 5.**
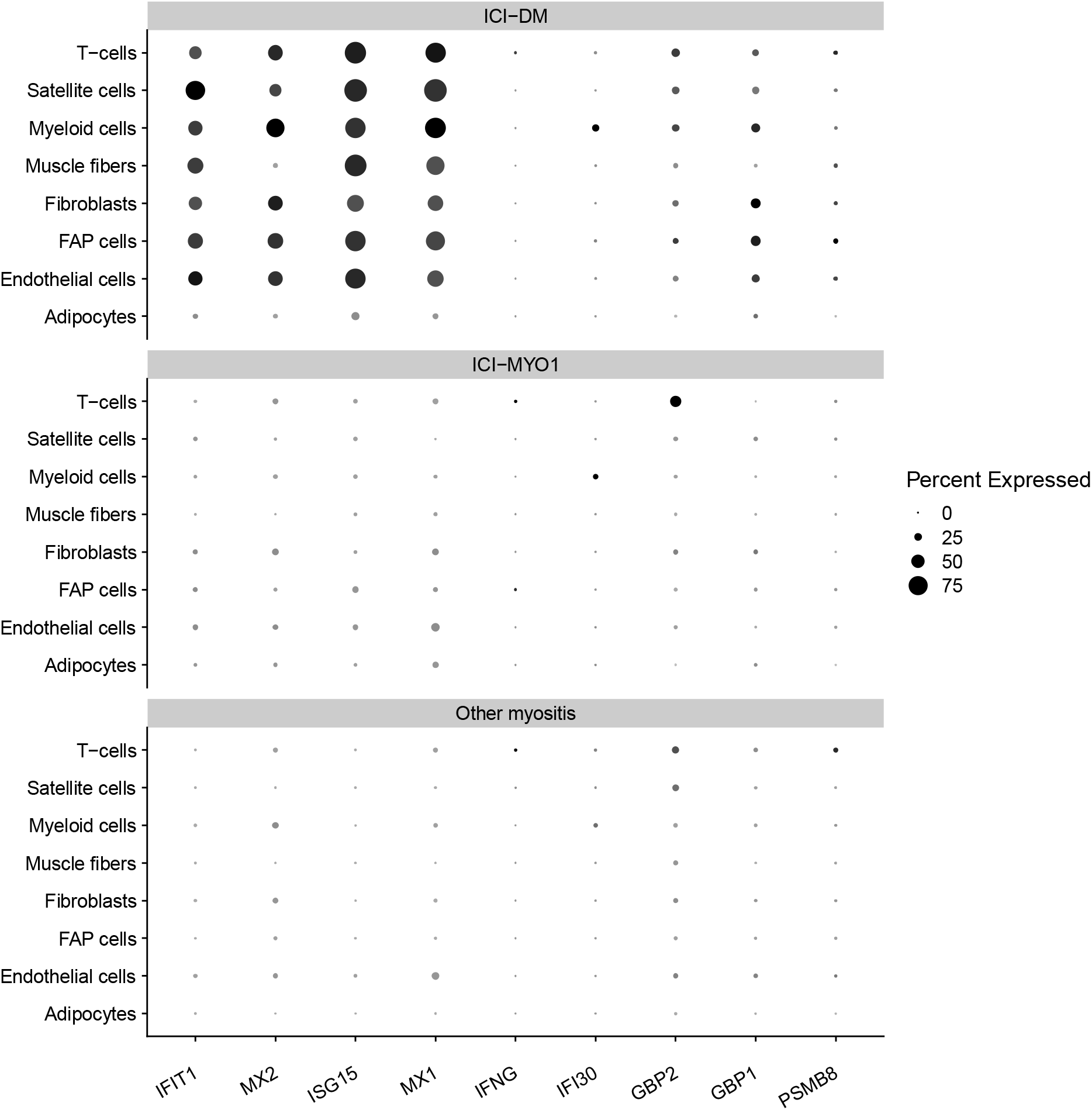
Percent of cells expressing genes related to the type 1 (IFIT1, MX2, ISG15, MX1) and type 2 (IFNG, IFI30, GBP2, GBP1, PSMB8) interferon pathway in muscle biopsies from clusters ICI-DM, and ICI-MYO1 of immune checkpoint-induced myopathy compared with a representative selection of patients with other types of inflammatory myopathy (4 dermatomyositis, 3 antisynthetase syndrome, 6 immune-mediated necrotizing myositis, and 2 inclusion body myositis).

### IL6 pathway genes are specifically up-regulated in all patients with ICI-myositis

Pathway analysis showed overexpression of signaling by interleukins (Figure 4). Of all the interleukins, genes associated with the IL6 pathway were the most specifically overexpressed in muscle biopsies from patients with ICI-MYO1, ICI-MYO2, and ICI-DM. These included genes encoding the IL6 receptor (IL6R), STAT3, and TYK2 (Figure 3–4, Supplementary Figure 5-7, Supplementary Table 3). Furthermore, IL6 expression itself was positively correlated with the level of expression of canonical inflammatory T-cell markers including CD4 and CD8 as well as the macrophage markers CD14 and CD68 (Supplementary Figure 8).

The gene encoding CEBPB, which binds to regulatory regions of several acute-phase and cytokines genes, including IL6, was also specifically overexpressed in all patients with ICI-myositis (ICI-MYO1, ICI-MYO2, and ICI-DM). Also, genes implicated in the suppression of the IL6 pathway, like SOCS3,[16] were more activated in ICI-MYO1 than in ICI-MYO2 (Figure 3, Supplementary Figure 6, Supplementary Table 3).

The expression levels of JUN, FOS, and EGR1 correlated with the expression of IL6 (Supplementary Figure 9) and were overexpressed in all three ICI-myositis clusters (ICI-MYO1, ICI-MYO2, and ICI-DM) (Supplementary Figure 10). In general, the overexpression of these genes was more intense in ICI-MYO1 and ICI-DM than in ICI-MYO2 (Supplementary Figure 10). Both EGR1, and the members of the transcription factor complex Activator Protein-1 JUN and FOS, are regulators of the IL6 pathway.[17–20]

Single-nuclei data confirmed that the IL6 pathway and its regulators were robustly overexpressed both in ICI-MYO1 and ICI-DM (Figure 6, Supplementary Tables 4-5). This study also showed that IL6R was expressed primarily in macrophages (Figure 6), although its expression was at levels too low to detect differences between groups (Supplementary Tables 4-5).

**Figure 6.**
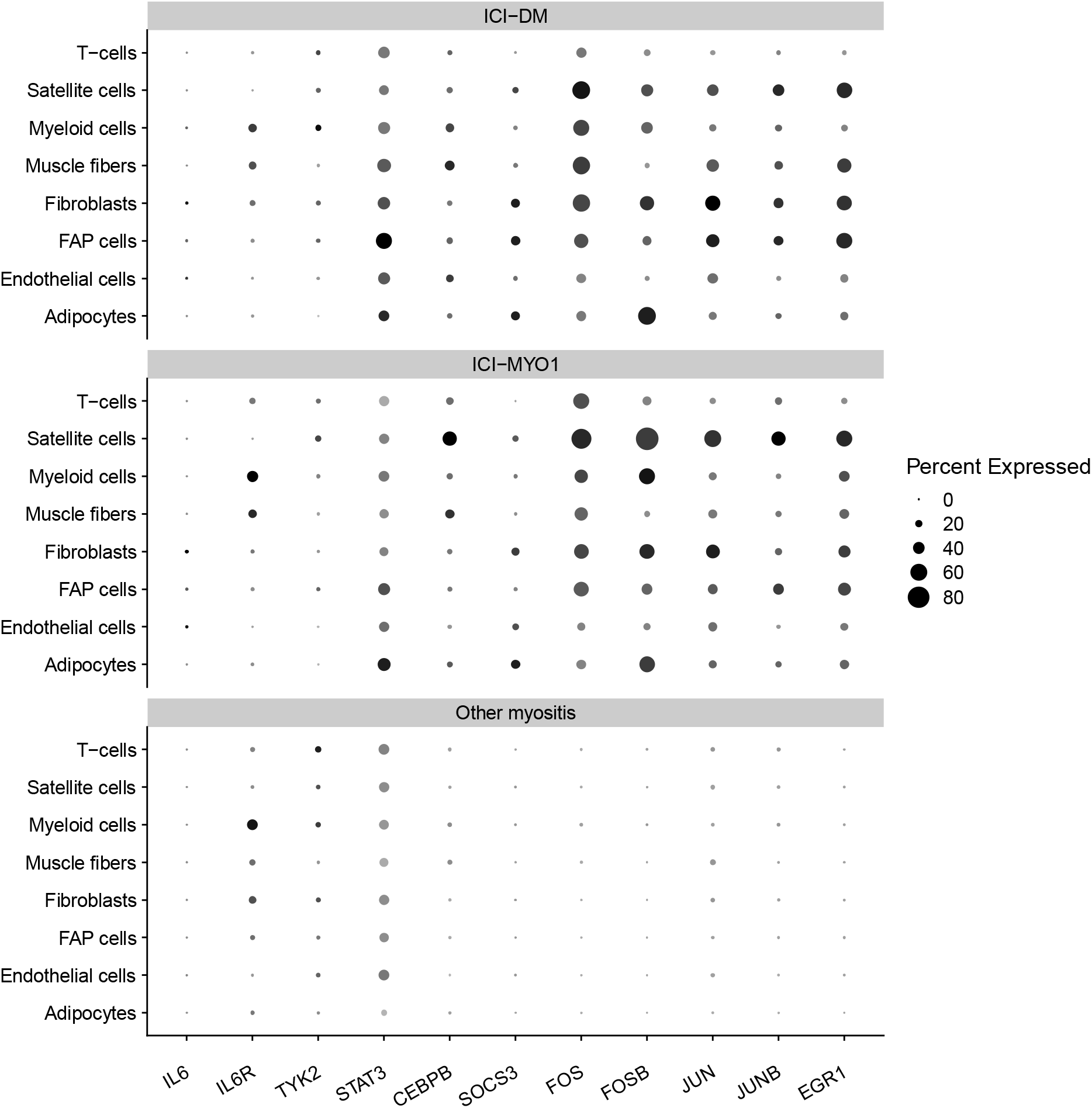
Percent of cells expressing genes related to the IL6 pathway in muscle biopsies from immune checkpoint-induced myopathy from clusters ICI-DM, and ICI-MYO1 compared with a representative selection of patients with other types of inflammatory myopathy (4 dermatomyositis, 3 antisynthetase syndrome, 6 immune-mediated necrotizing myositis, and 2 inclusion body myositis).

#### Type 1 interferon-inducible genes are overexpressed in ICI-DM

Compared to patients in the ICI-MYO1 or ICI-MYO2 clusters, the three patients within the ICI-DM cluster had a marked elevation of type I interferon-stimulated genes, as seen in DM patients (Figure 3–4, Supplementary Figure 11-12). In these ICI-DM patients, IFNB1 gene expression was detectable (Supplementary Figure 13). In contrast, other type-I interferon genes were either undetectable or present at lower levels, as in muscle biopsies from patients with DM (Supplementary Figure 13, Supplementary Table 3).

Single-nuclei RNAseq verified a robust activation of the type 1 interferon pathway affecting all the different cell types of patients with ICI-DM (Figure 5, Supplementary Tables 4-5)

#### Other transcriptomic features of ICI-myositis compared to DM, AS, IMNM, IBM, and healthy comparator muscle

Compared to control muscle tissue, muscle biopsies from patients with ICI-MYO1 and ICI-DM had increased expression of genes associated with macrophages (CD14, and CD68) and T-cells (CD3E, CD4, CD8A, PRF1, GZMA, GZMB) along with reductions in the expression of skeletal muscle structural genes (MYH1, ACTA1). Genes upregulated during muscle regeneration (NCAM1, MYH3) were elevated in both ICI-MYO1 and ICI-DM. Genes associated with oxidative phosphorylation and mitochondrial genes had decreased expression; this was more pronounced in ICI-MYO1 and ICI-DM than in ICI-MYO2. The expression of immunoglobulin genes was increased only in ICI-MYO1; levels of these genes were similar to those in DM and AS, but lower than in IBM (Supplementary Figure 14-17, Supplementary Table 3).

Multiple TNF receptors and their ligands were exclusively overexpressed in patients with ICI, including those signaling TNFa (Supplementary Figure 18-20, Supplementary Table 3). Accordingly, the TNFa pathway was overexpressed in patients with ICI-MYO1 and ICI-DM, similar to other types of inflammatory myopathy.

Also, vascular adhesion molecules like VCAM1 and ICAM1 were overexpressed at similar levels to other types of inflammatory myopathy in ICI-DM and ICI-MYO1, but not in ICI-MYO2 (Supplementary Figure 21, Supplementary Table 3).

Finally, several checkpoint genes such as PDCD1, and CTLA4 were upregulated in ICI-MYO and ICI-DM, but not in ICI-MYO2 (Supplementary Figure 22).

#### Comparing transcriptomic profiles of ICI-myositis patients with and without ocular and cardiac involvement

We did not find any significant transcriptomic differences between ICI-myositis patients with and without diplopia or between ICI-myositis patients exposed to different types of ICI (PD-1 inhibitors, PD-L1 inhibitors, and co-treatment with CTLA4). However, patients with myocarditis were all part of cluster ICI-MYO1 and had higher levels of IFNG (log2FC 2.8, q-value 0.03), CD8A (log2FC 2.3, q-value 0.03), CD14 (log2FC 1.9, q-value 0.05), and vascular adhesion molecules (ICAM1 log2FC 1.3, q-value 0.05) (Figure 7, Supplementary Table 3).

**Figure 7.**
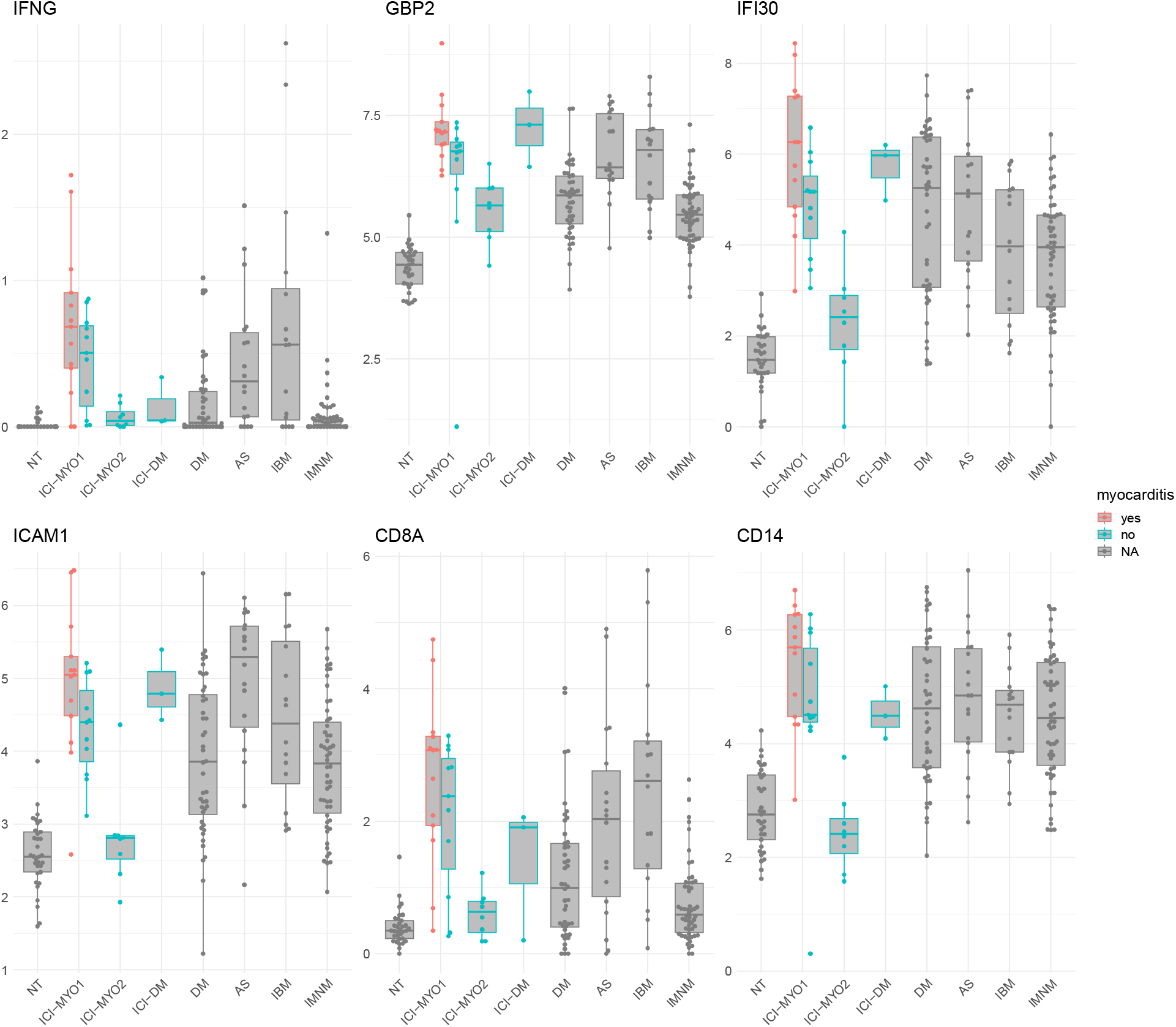
Expression levels (log2[TMM + 1]) of IFNG, IFNG-inducible genes, ICAM1, CD8A, and CD14 in patients according to the presence of myocarditis. NT: normal muscle; DM: dermatomyositis; AS: antisynthetase syndrome; IBM: inclusion body myositis; IMNM: immune-mediated necrotizing myopathy.

#### Tumors treated with ICI upregulate IFNγ-stimulated genes but not genes of the IL6 pathway

We were interested in whether ICI might cause upregulation of the IL6 pathway even in the absence of myositis. Unfortunately, muscle biopsies from patients who were treated with ICI but who did not develop myositis were not available. However, publicly available RNA sequencing data from melanomas before and after treatment with ICI (GEO: GSE91061) showed overexpression of IFNG (log2FC 1.2, q-value=0.008) and IFNG-stimulated genes after treatment with ICI (e.g. GBP2 log2FC 0.8, q-value=0.03). Unlike the muscle of patients with ICI-myositis, these tumors did not show a significant overactivation of the IL6 pathway after treatment with ICI (e.g. IL6R log2FC 0.008, q-value 0.8, Supplementary Figure 23).

## Discussion

In this study, we identified three transcriptomically distinct clusters of ICI-myositis patients and showed that each cluster includes patients with unique clinical features. At the transcriptomic level, all three clusters were characterized by over-expression of the IL6 pathway. In contrast, only biopsies from the ICI-DM and ICI-MYO1 clusters had high expression of IFNγ-stimulated genes, whereas IFNB1 and IFNβ1-inducible genes were only highly expressed in biopsies from the ICI-DM cluster. From the clinical perspective, only patients in the ICI-MYO1 cluster developed myocarditis and only patients in the ICIDM cluster had DM rashes.

The pathogenesis of autoimmune adverse events in the context of immune checkpoint inhibitor treatment is still not completely understood. One hypothesis is that immune-related adverse events may be caused by a sudden and intense activation of already-existing autoimmunity. In this study, we provide two pieces of information supporting this theory. First, the autoantibodies present at the time of ICI-myositis were also present before ICI treatment in all three patients for whom pre-treatment sera were available. Second, patients with ICI-DM recapitulated the key transcriptomic features of patients with dermatomyositis and had the characteristic autoantibodies of patients with paraneoplastic dermatomyositis (anti-TIF1γ).

Of note, our group has previously shown that in patients with thymoma and ICI-myositis, anti-AChR autoantibodies are detectable prior to the start of ICI therapy.[21] Interestingly, the most common autoantibodies detected in the patients of our study, anti-AChR, and anti-TIF1g autoantibodies, are also common autoantibodies associated with cancer in patients with myasthenia gravis and myositis,[22, 23] suggesting the possibility that the preexisting autoimmune phenomenon that is activated by ICI may have been directly induced by the tumor itself prior to ICI therapy.

ICI-myositis patients in cluster ICI-MYO1 had frequent myocarditis and autoantibodies targeting the neuromuscular junction (anti-AChR autoantibodies) or skeletal muscle (anti-striated muscle autoantibodies). These features are uncommon in patients with other types of inflammatory myopathy and have also been described in cases of myasthenia gravis not related to ICI.[24] Moreover, many of these patients had clinical features typical of myasthenia gravis, such as diplopia. Autoantibodies in myasthenia gravis bind to the surface of the muscle (postsynaptic receptors of the neuromuscular junction), and thus, these autoantibodies bound to the muscle fibers may have a role attracting to the muscle autoreactive T cells activated by the ICI. Supporting this theory, we have found that patients that develop myocarditis are the ones that have the most active expression of IFNG-inducible genes, T-cell markers, and vascular adhesion molecules.

Our findings may have potential therapeutic implications. Corticosteroids as well as other agents such as IVIG[25] and abatacept[26] have been used to treat ICI-myositis. However, it remains unknown how effective they are. Furthermore, in the case of abatacept, there is reason to be concerned that its binding to CD80/86 may counteract the beneficial effects of ICI therapy on the tumor. In this study, we identified the IL6 pathway as being specifically overexpressed in ICI-myositis and not elevated in tumors treated with ICI. Supporting our findings, the IL6 pathway has been shown to be elevated in the affected tissue of patients with colitis induced by ICI.[27] Furthermore, tocilizumab, a blocker of the IL6 receptor, has been reported as a potentially effective treatment for various ICI-triggered immune adverse events.[27, 28] Interestingly, it was reported that blocking this pathway may have beneficial effects on the antineoplastic effects of ICI.[27] Thus, it is possible that the overexpression of the IL6 pathway is a general phenomenon not restricted to muscle in ICI autoimmune adverse events, and that treatment with tocilizumab may be useful not only to treat or prevent the adverse event but also to improve tumor prognosis. The fact that type 2 IFN is overexpressed in all ICI-MYO, and type 1 IFN in ICIDM, also suggests that targeting the JAK-STAT pathway may be useful in patients with ICI-myositis, but risks negatively impact the antineoplastic effect of the drug. Notwithstanding this, given the quick effect of JAK-STAT inhibitors, it may be reasonable to use them in patients with severe cases of ICI-myositis.

This study has several limitations. First, most patients did not have sera available before the start of ICI. Furthermore, the techniques use to assess serologic profiles varied between the different participating centers. Also, there was heterogeneity among patients both in the type of tumors they had and the type of ICI they received. Furthermore, given the severity of their clinical manifestations, half of the patients were treated with corticosteroids before the biopsy was performed (median duration 6 days), which may have reduced the sensitivity of our analyses. These caveats notwithstanding, the lack of heterogeneity in the relatively large number of samples studied may also make our conclusions more generalizable. Finally, compared to bulk RNAseq, single-nuclei RNA sequencing may not detect genes expressed at low levels and this limited our ability to explore some of the affected pathways in greater detail.

Despite these limitations, this study reveals the existence of three transcriptomically distinct types of ICI-myositis and demonstrates that each type has distinct clinical features. We also demonstrate that the IL6 pathway is activated in all three types of ICI-myositis but not in other types of ICI-naive myositis such as DM, AS, IMNM, or IBM. Based on the evidence provided by this study and previously published studies, we propose that targeting the IL6 pathway may be therapeutically useful in all three types of ICI-myositis patients.

## Acknowledgments

We would like to acknowledge Helena Verdaguer for her kind contribution to this research. This study was funded, in part, by the Intramural Research Programs of the National Institute of Arthritis and Musculoskeletal and Skin Diseases (NIAMS), the National Cancer Institute (NCI), the Center for Cancer Research of the National Institutes of Health, and through a Cooperative Research and Development Agreement between the NCI and EMD Serono Research & Development Institute, Inc., Billerica, MA, USA, an affiliate of Merck KGaA (CrossRef Funder ID:10.13039/100004755), as part of an alliance between Merck and Pfizer. This work was also supported by the Peter Buck and the Huayi and Siuling Zhang Discovery Fund.

