## Supplementary Figures for "Transcriptomic profiling reveals distinct subsets of immune checkpoint inhibitor-induced myositis"

**Supplementary Figure 1.** Representative genes for each single-nuclei RNA sequencing cluster.

**Supplementary Figure 2.** Expression (log2[TMM+1]) of predominantly type1 interferon-stimulated genes (ISG15, and MX1), and predominantly type 2 interferon-stimulated genes (GBP2, and IFI30) in differentiating human skeletal muscle myoblasts treated with IFNA2a, IFNB1, and IFNG at two different doses each (100U, and 1000U).

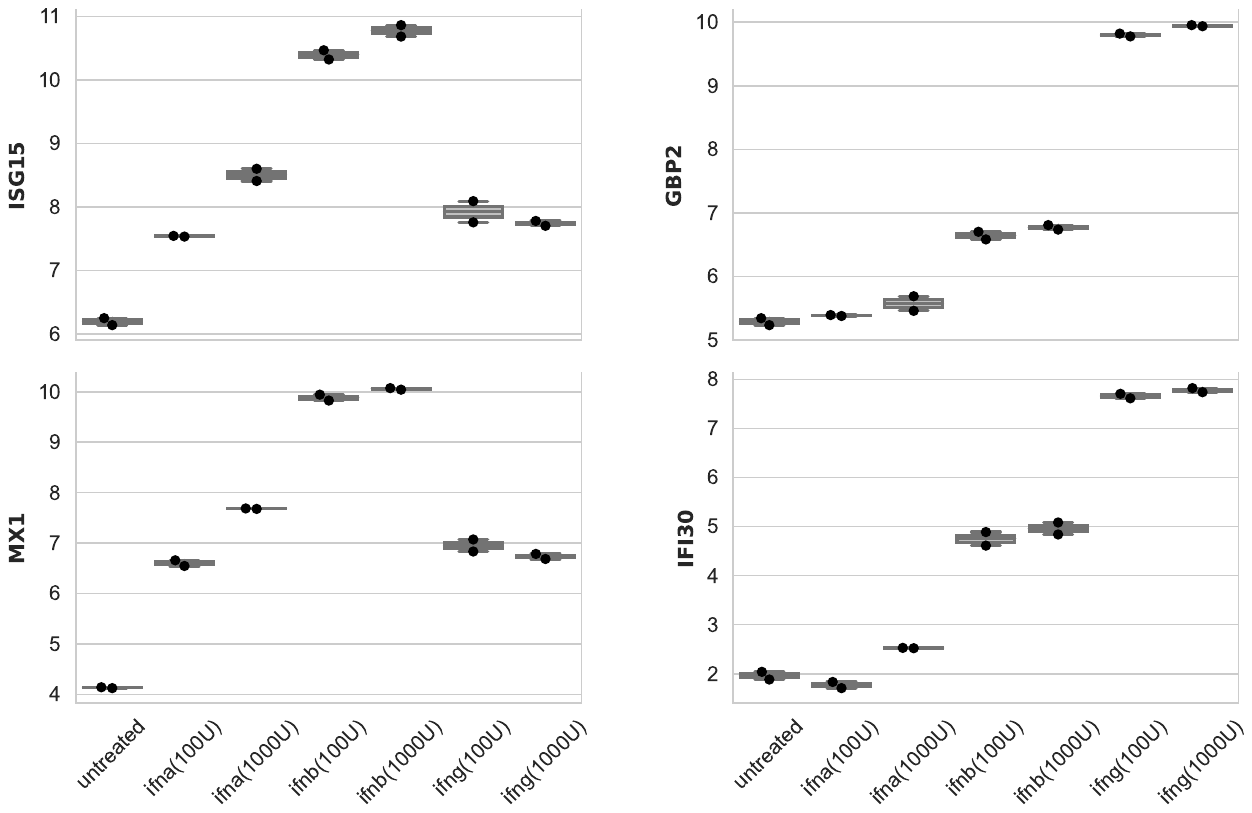

**Supplementary Figure 3.** Expression (log2[TMM+1]) of IFNG and representative IFNG-stimulated genes.

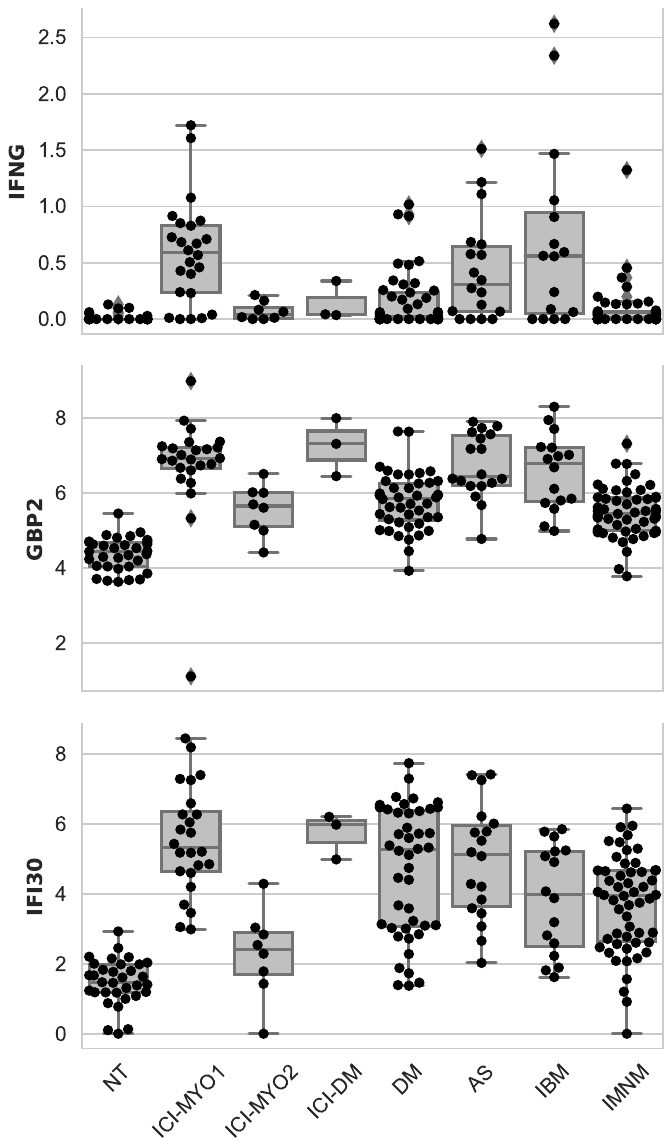

NT: normal muscle; ICI-DM: immune checkpoint-induced dermatomyositis; DM: dermatomyositis; AS: antisynthetase syndrome; IBM: inclusion body myositis; IMNM: immune-mediated necrotizing myopathy.

**Supplementary Figure 4.** Gene Set Enrichment Analysis of the interferon-gamma response (left) in immune checkpoint-induced myopathy patients compared to normal muscle (p-value 0.04). Fifty genes with the highest signal-to-noise ratio in this pathway (red high, blue low).

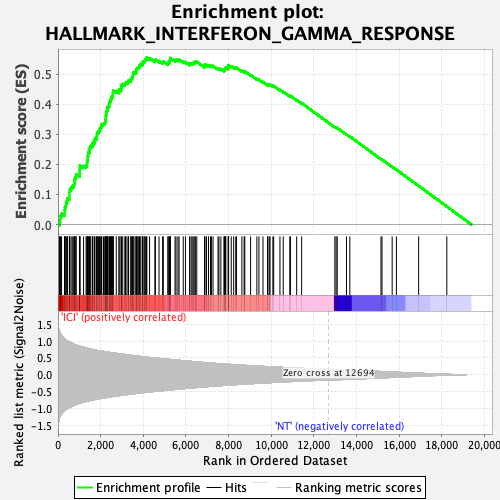

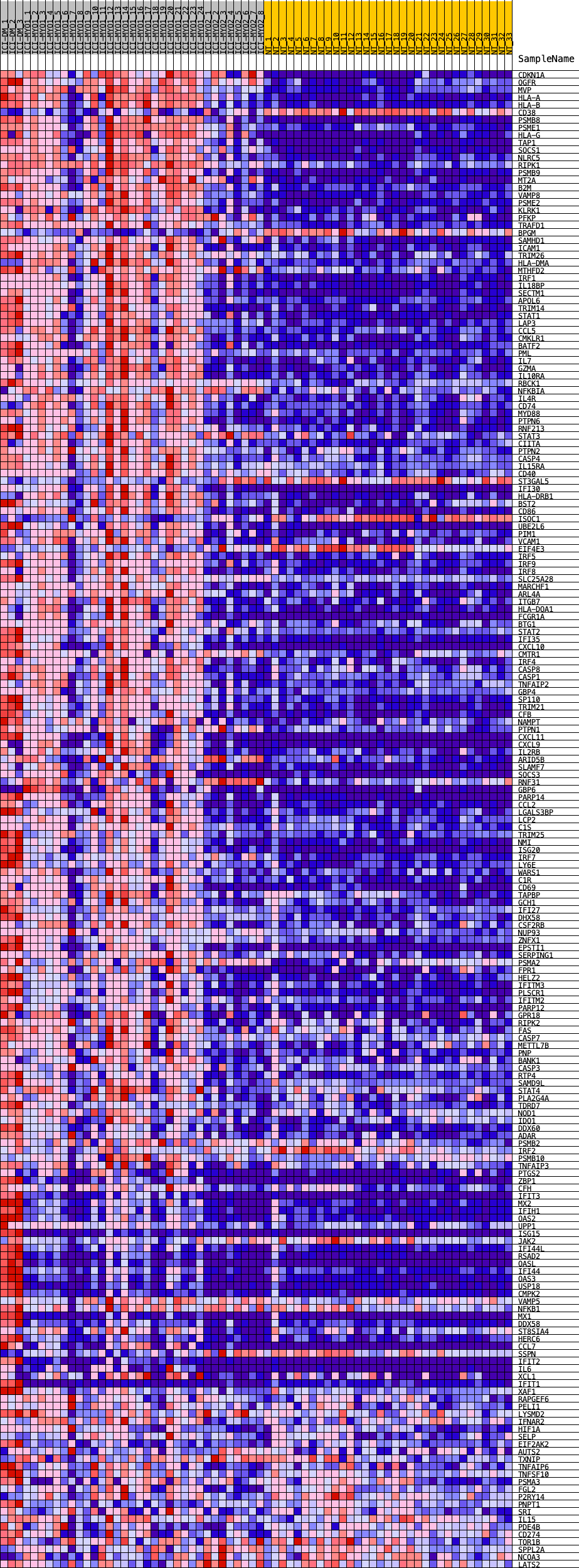

**Supplementary Figure 5.** Expression (average z-score of log2[TMM+1]) of cytokines and cytokine receptors in patients with ICI-induced myopathy (ICI-MYO1, ICI-MYO2, and ICI-DM) and in the comparator muscles biopsies.

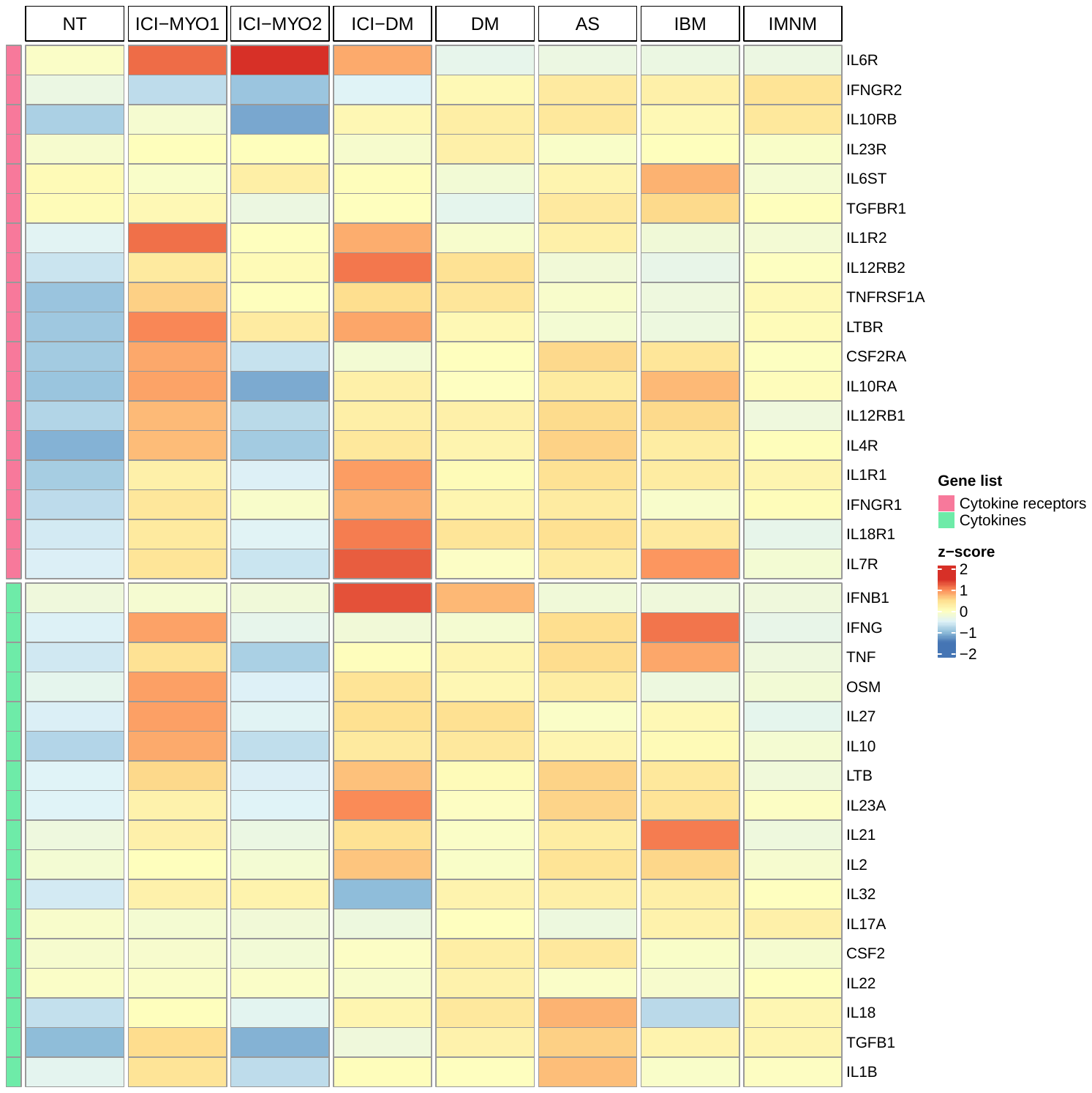

NT: normal muscle; DM: dermatomyositis; AS: antisynthetase syndrome; IBM: inclusion body myositis; IMNM: immune-mediated necrotizing myopathy.

**Supplementary Figure 6.** Expression (log2[TMM+1]) of representative genes from the IL6 pathway.

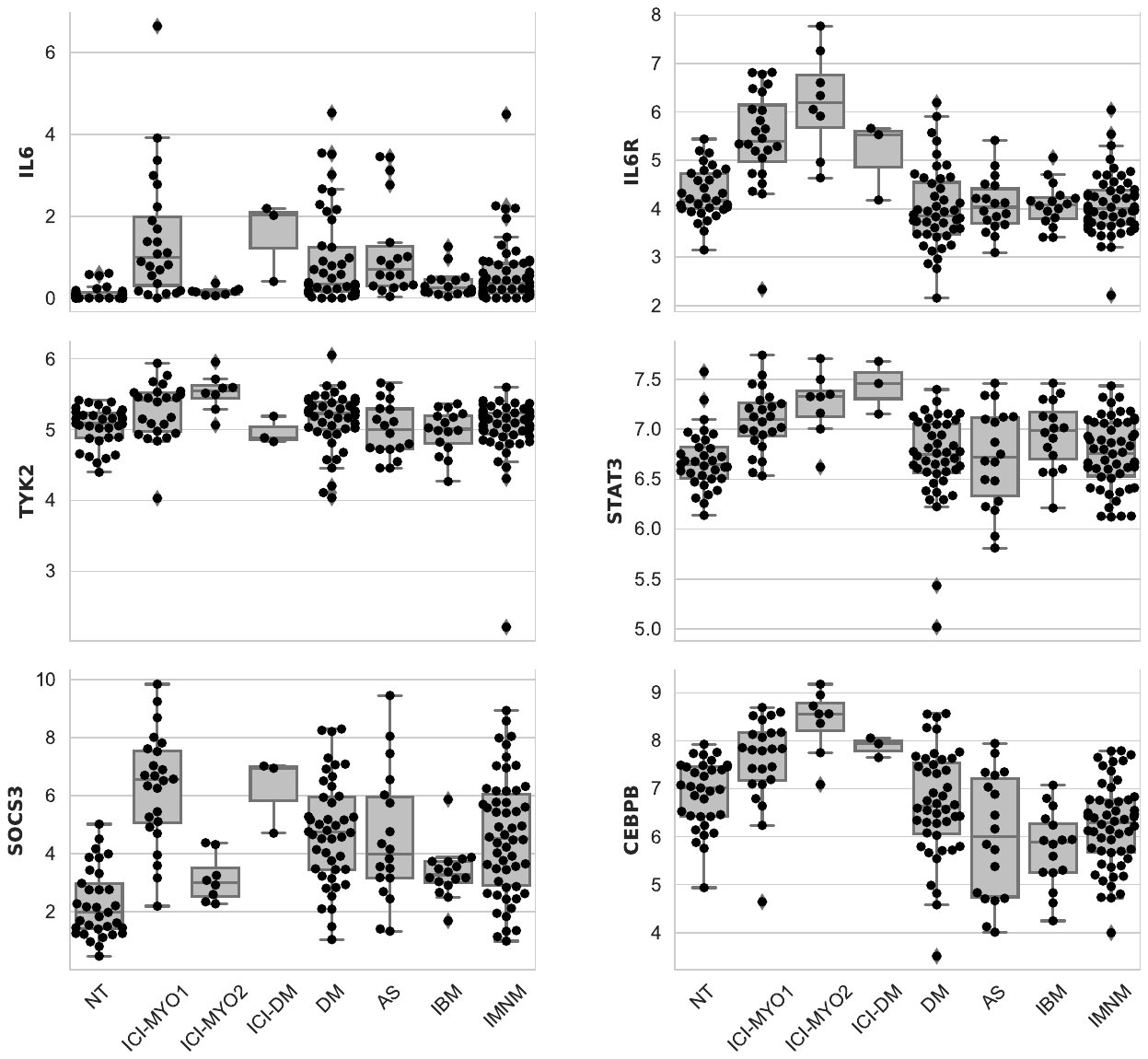

NT: normal muscle; ICI-DM: immune checkpoint-induced dermatomyositis; DM: dermatomyositis; AS: antisynthetase syndrome; IBM: inclusion body myositis; IMNM: immune-mediated necrotizing myopathy.

**Supplementary Figure 7.** Gene Set Enrichment Analysis of the IL6-JAK-STAT3 pathway (left) in immune checkpoint-induced myopathy patients compared to normal muscle (p-value 0.01). Fifty genes with the highest signal-to-noise ratio in this pathway (red high, blue low).

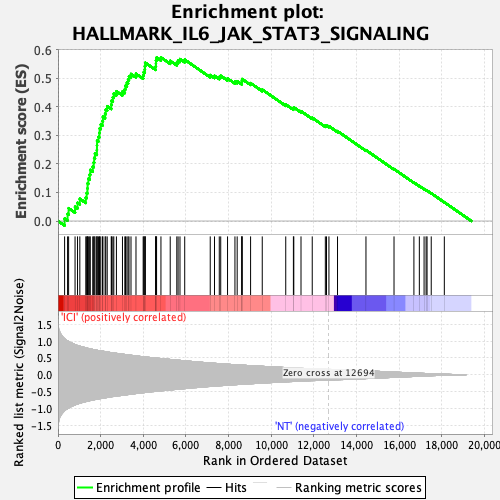

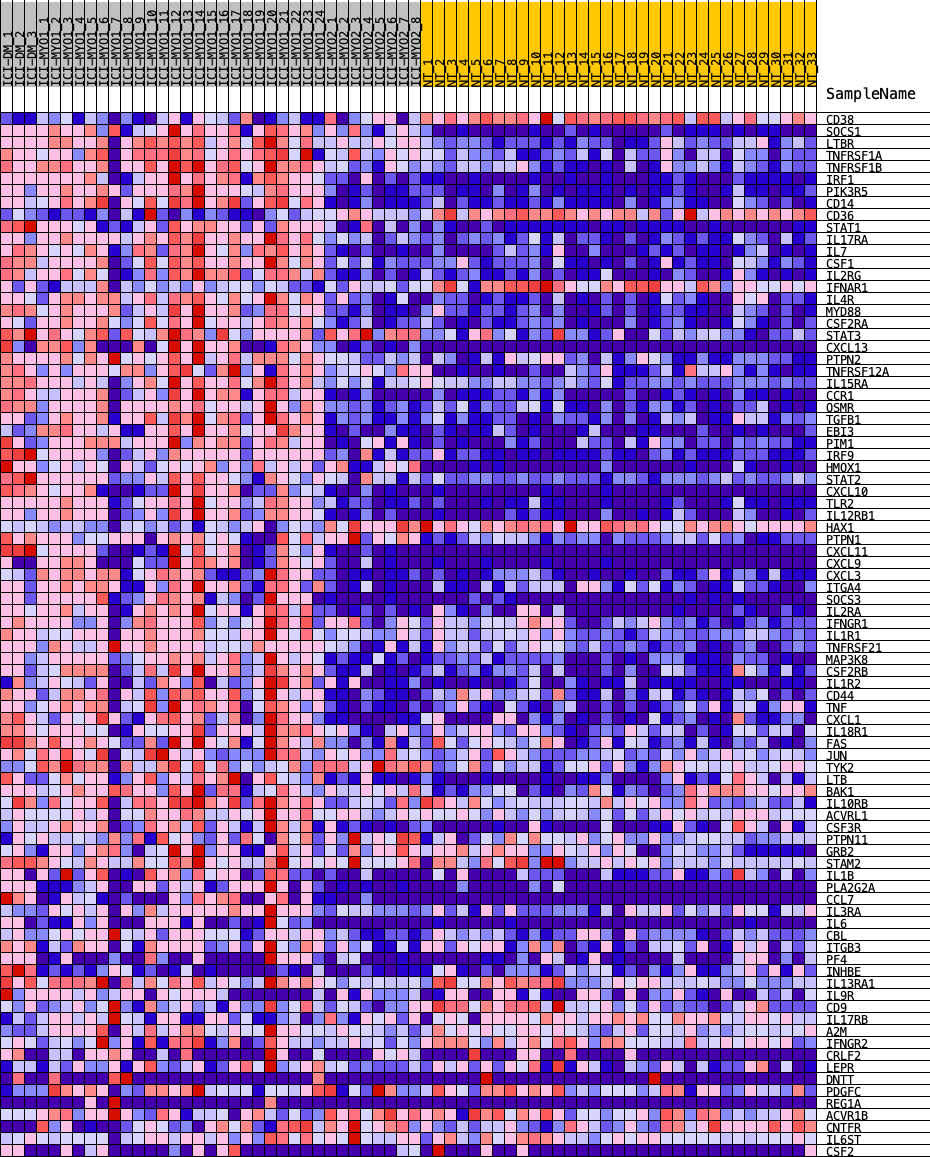

**Supplementary Figure 8.** Correlation of CD4, CD8A, CD14, and CD68 with IL6 in the different clusters of patients with immune checkpoint-induced myopathy (ICI-MYO1, ICI-MYO2, and ICI-DM)

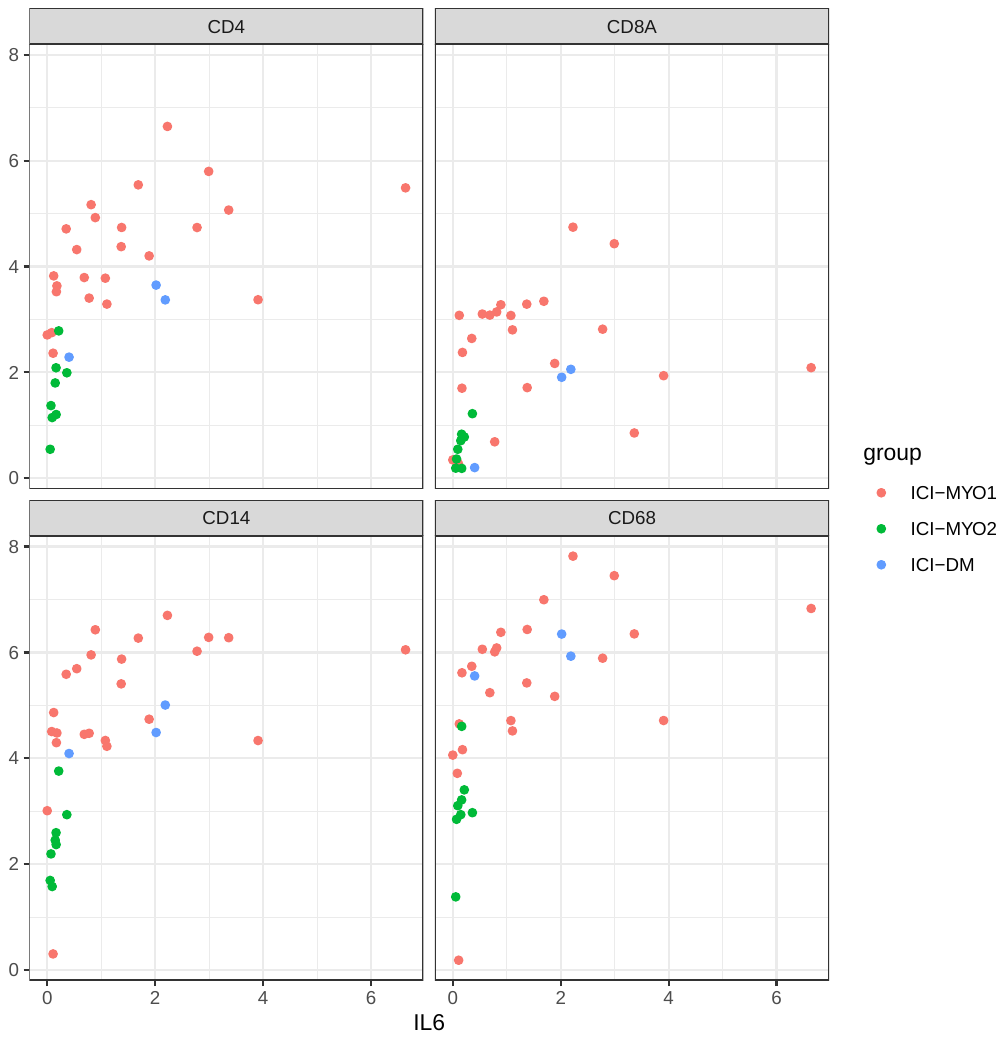

**Supplementary Figure 9.** Correlation of IL6 with EGR1, SOCS3, and members of the protein families FOS and JUN in the different clusters of patients with immune checkpoint-induced myopathy (ICI-MYO1, ICI-MYO2, and ICI-DM)

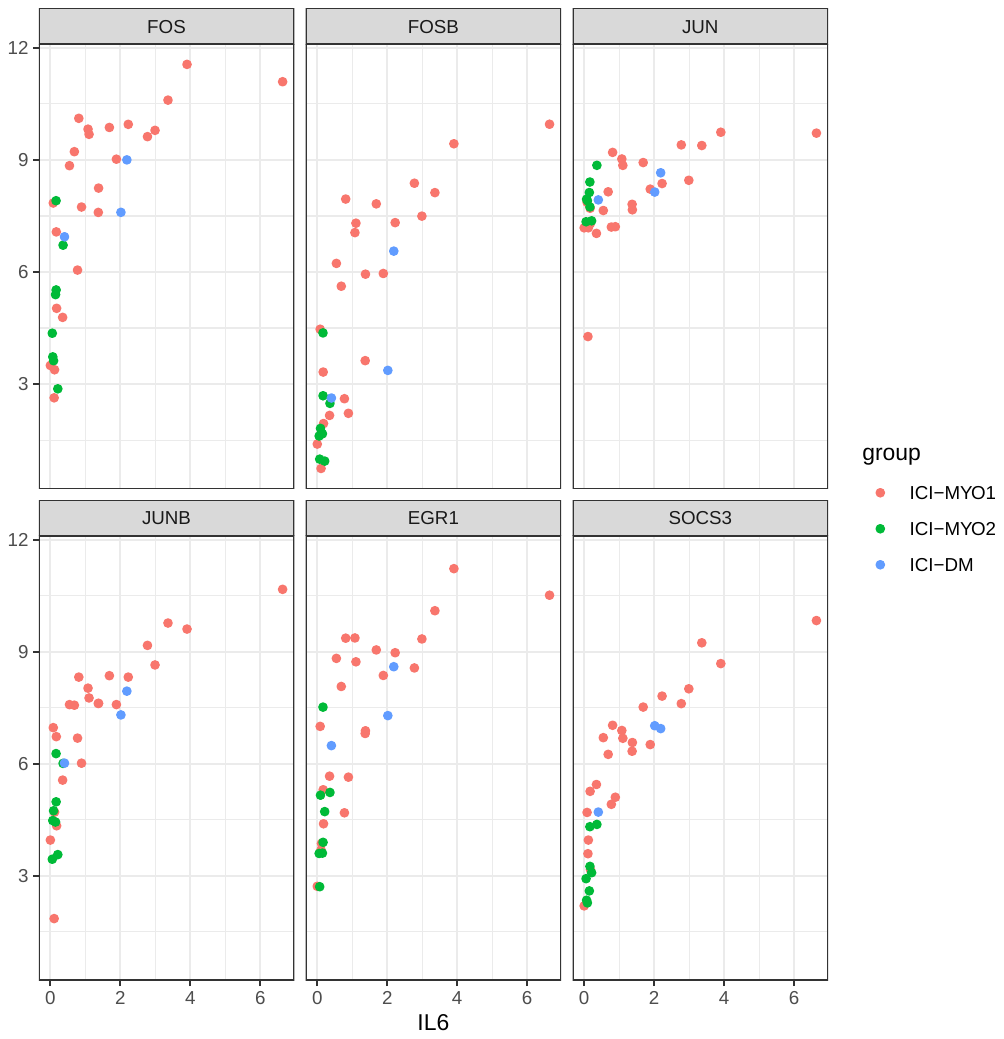

**Supplementary Figure 10.** Expression (log2[TMM+1]) of EGR1 and members of the FOS and JUN family of proteins.

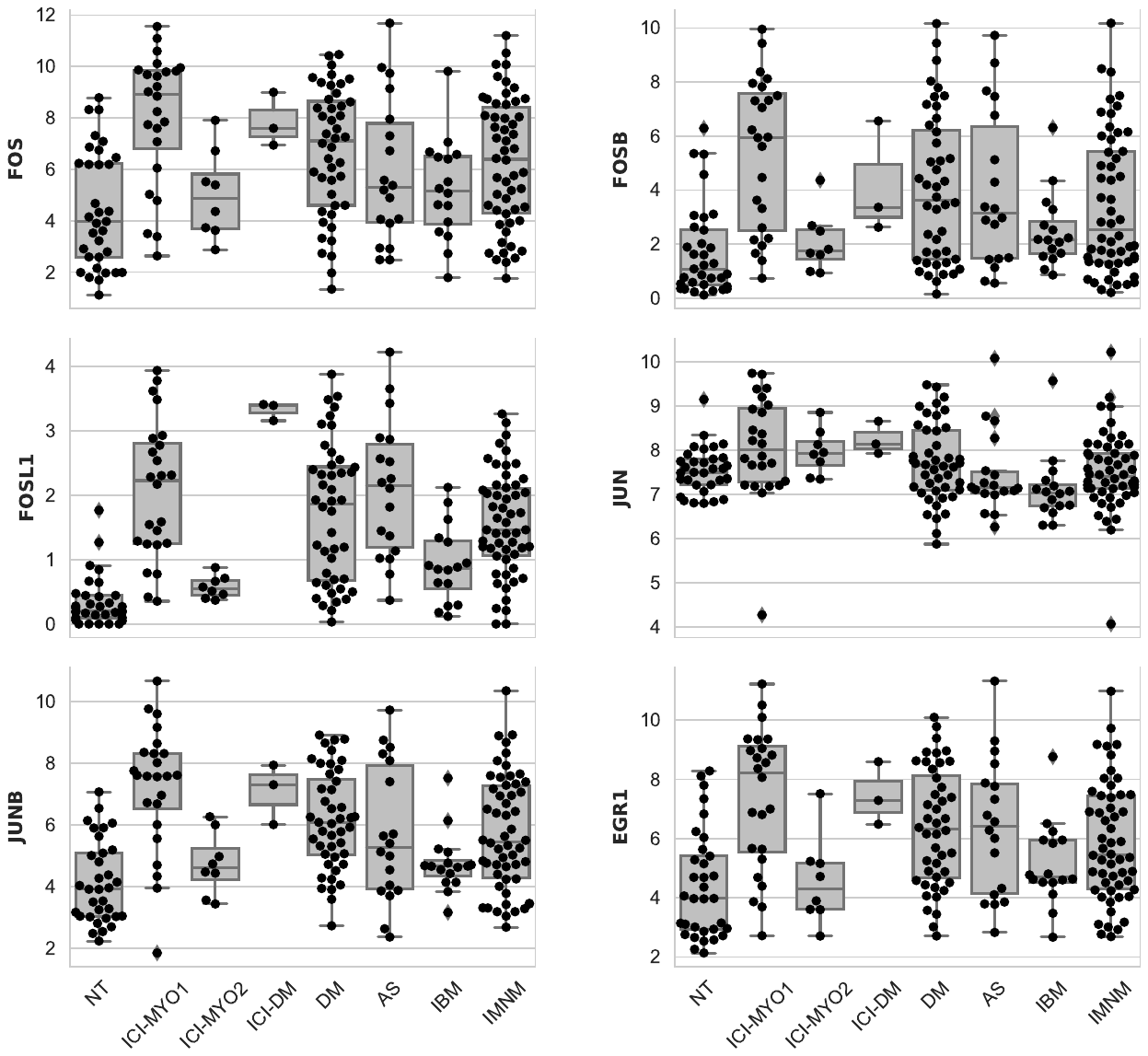

NT: normal muscle; ICI-DM: immune checkpoint-induced dermatomyositis; DM: dermatomyositis; AS: antisynthetase syndrome; IBM: inclusion body myositis; IMNM: immune-mediated necrotizing myopathy.

**Figure 11.** Expression levels (log2[TMM + 1]) of IFNB1, and interferon type I inducible genes ISG15 and MX1 in muscle. In patients with immune-checkpoint inhibitor-induced dermatomyositis (ICI-DM), IFNB1 and IFN type I inducible genes are overexpressed, similar to patients with non-ICI dermatomyositis.

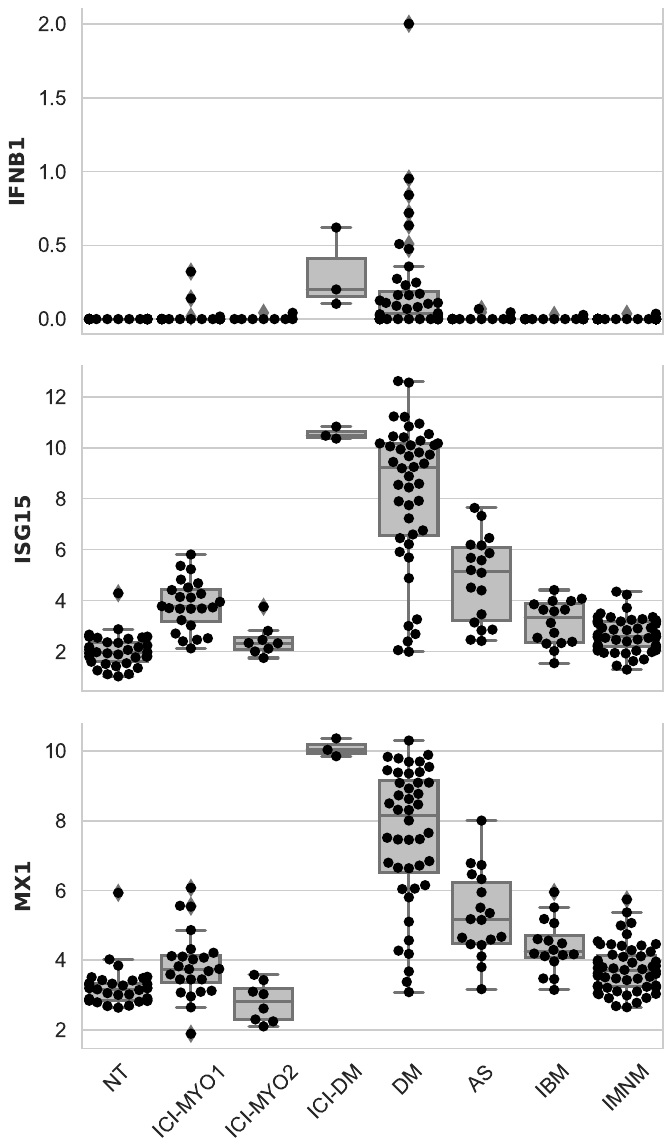

NT: normal muscle; ICI-DM: immune checkpoint-induced dermatomyositis; DM: dermatomyositis; AS: antisynthetase syndrome; IBM: inclusion body myositis; IMNM: immune-mediated necrotizing myopathy.

**Supplementary Figure 12.** Gene Set Enrichment Analysis of the type 1 interferon pathway (left) in immune checkpoint-induced dermatomyositis patients compared to normal muscle (p-value < 0.001). Fifty genes with the highest signal-to-noise ratio in this pathway (red high, blue low).

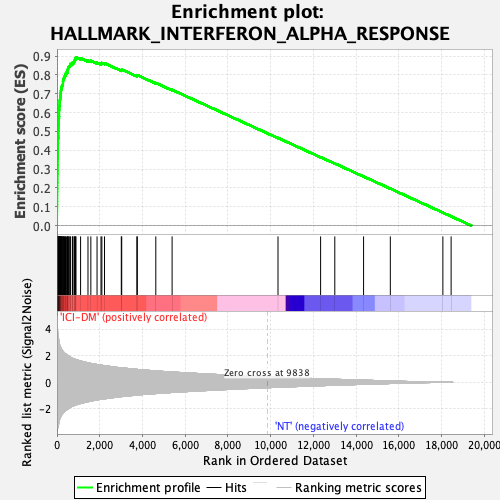

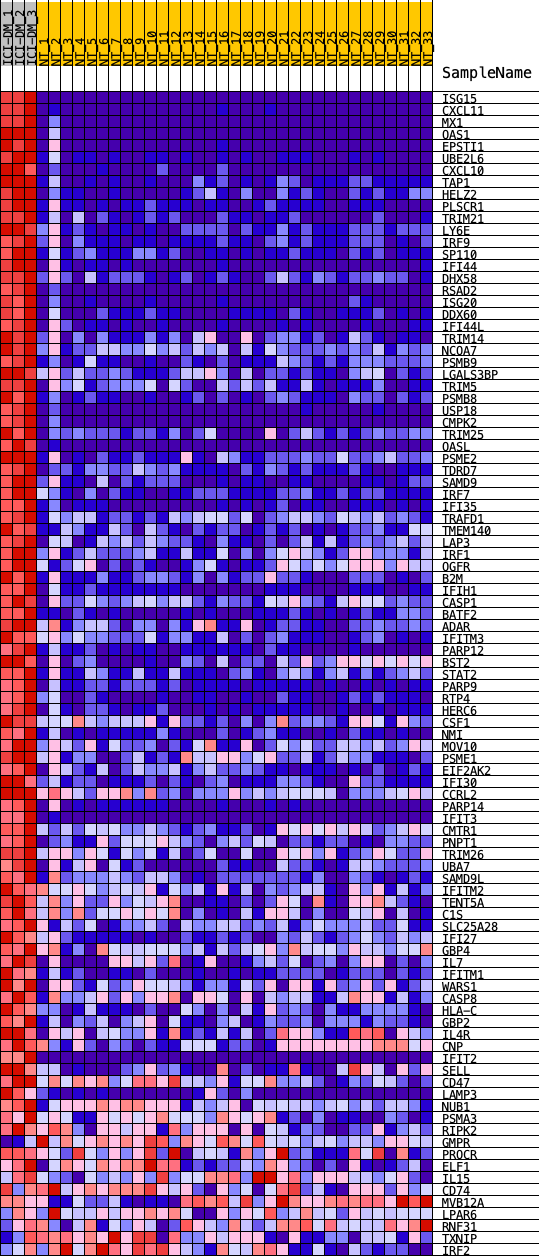

**Supplementary Figure 13.** Average expression levels (average and 95% confidence interval of log2[TMM + 1]) of type I interferon genes in muscle. Patients with immune checkpoint dermatomyositis have levels of IFNB1 similar to patients with dermatomyositis.

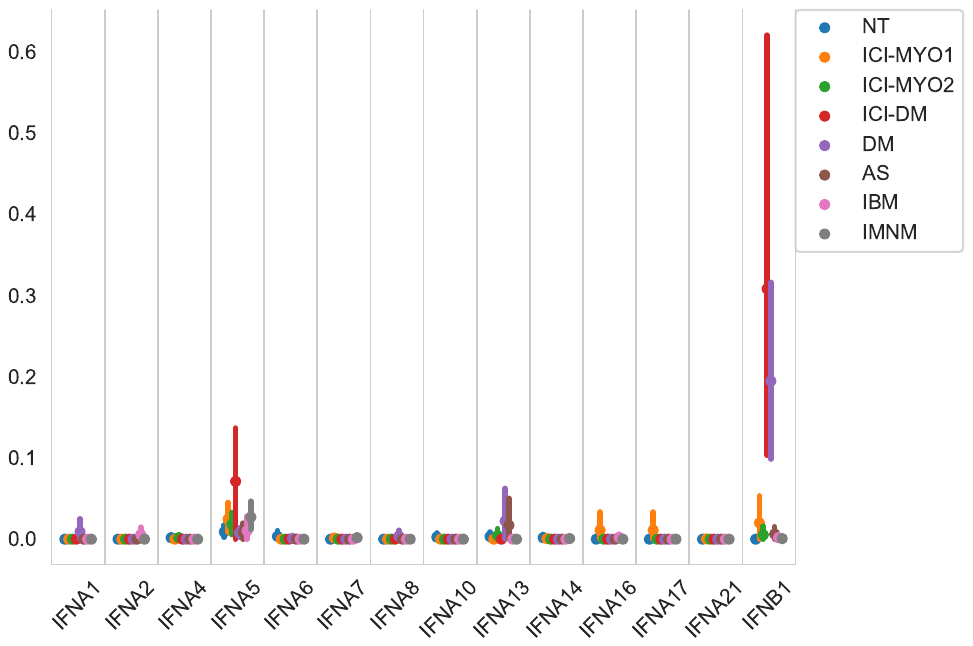

NT: normal muscle; DM: dermatomyositis; AS: antisynthetase syndrome; IBM: inclusion body myositis; IMNM: immune-mediated necrotizing myopathy

**Supplementary Figure 14.** Expression (log2[TMM+1]) of representative gene markers associated with B cells, macrophages, and T-cells.

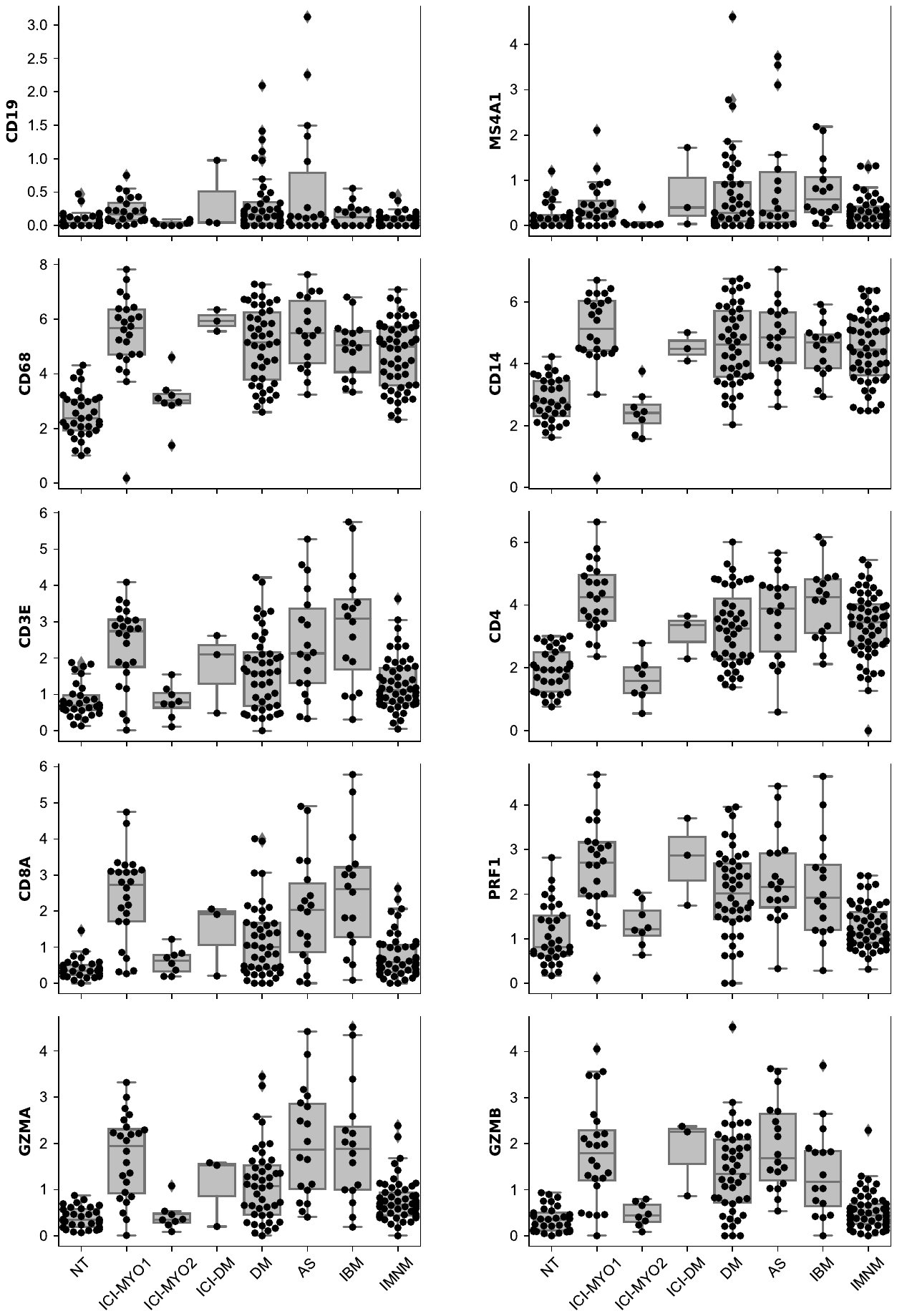

NT: normal muscle; DM: dermatomyositis; AS: antisynthetase syndrome; IBM: inclusion body myositis; IMNM: immune-mediated necrotizing myopathy.

**Supplementary Figure 15.** Expression (log2[TMM+1]) of representative gene markers associated with muscle regeneration, adult skeletal muscle, oxidative phosphorylation, and mitochondrial function.

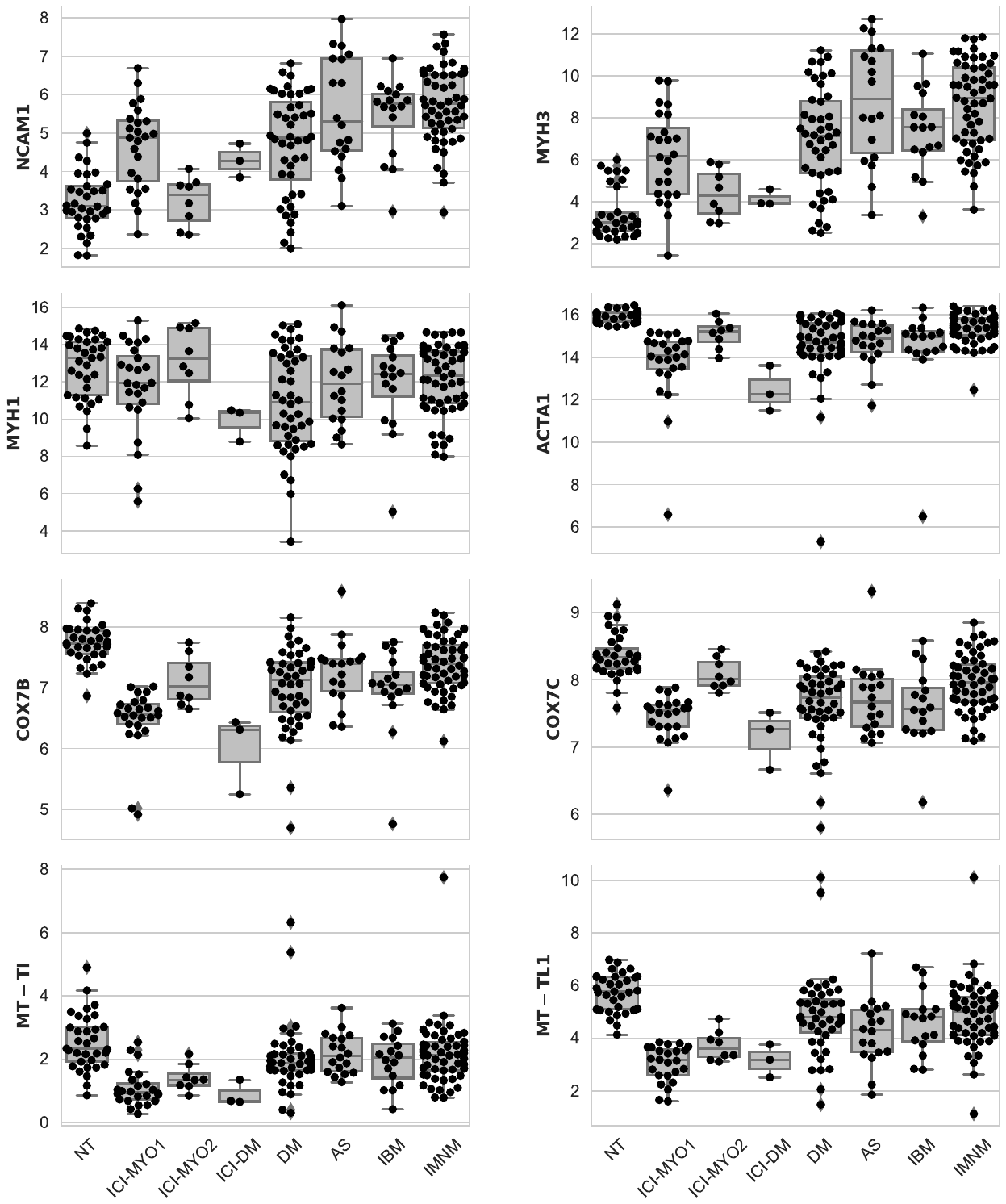

NT: normal muscle; DM: dermatomyositis; AS: antisynthetase syndrome; IBM: inclusion body myositis; IMNM: immune-mediated necrotizing myopathy.

**Supplementary Figure 16.** Gene Set Enrichment Analysis of the oxidative phosporylation pathway (left) in immune checkpoint-induced myopathy patients compared to normal muscle (p-value 0.006). Fifty genes with the highest signal-to-noise ratio in this pathway (red high, blue low).

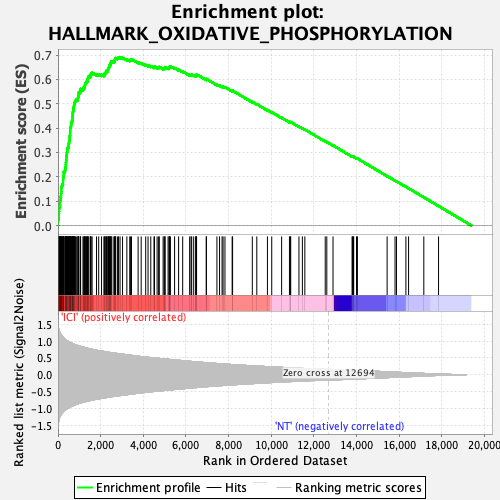

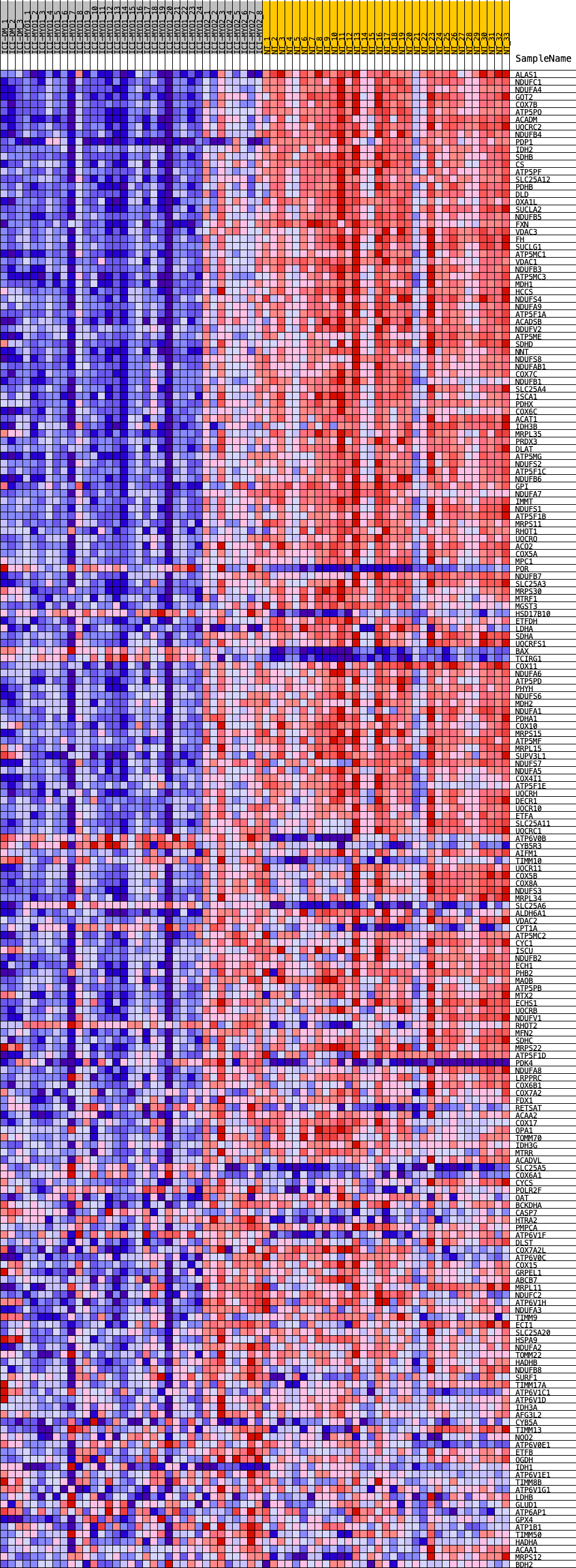

**Supplementary Figure 17.** Expression (log2[TMM+1]) of immunoglobulin genes.

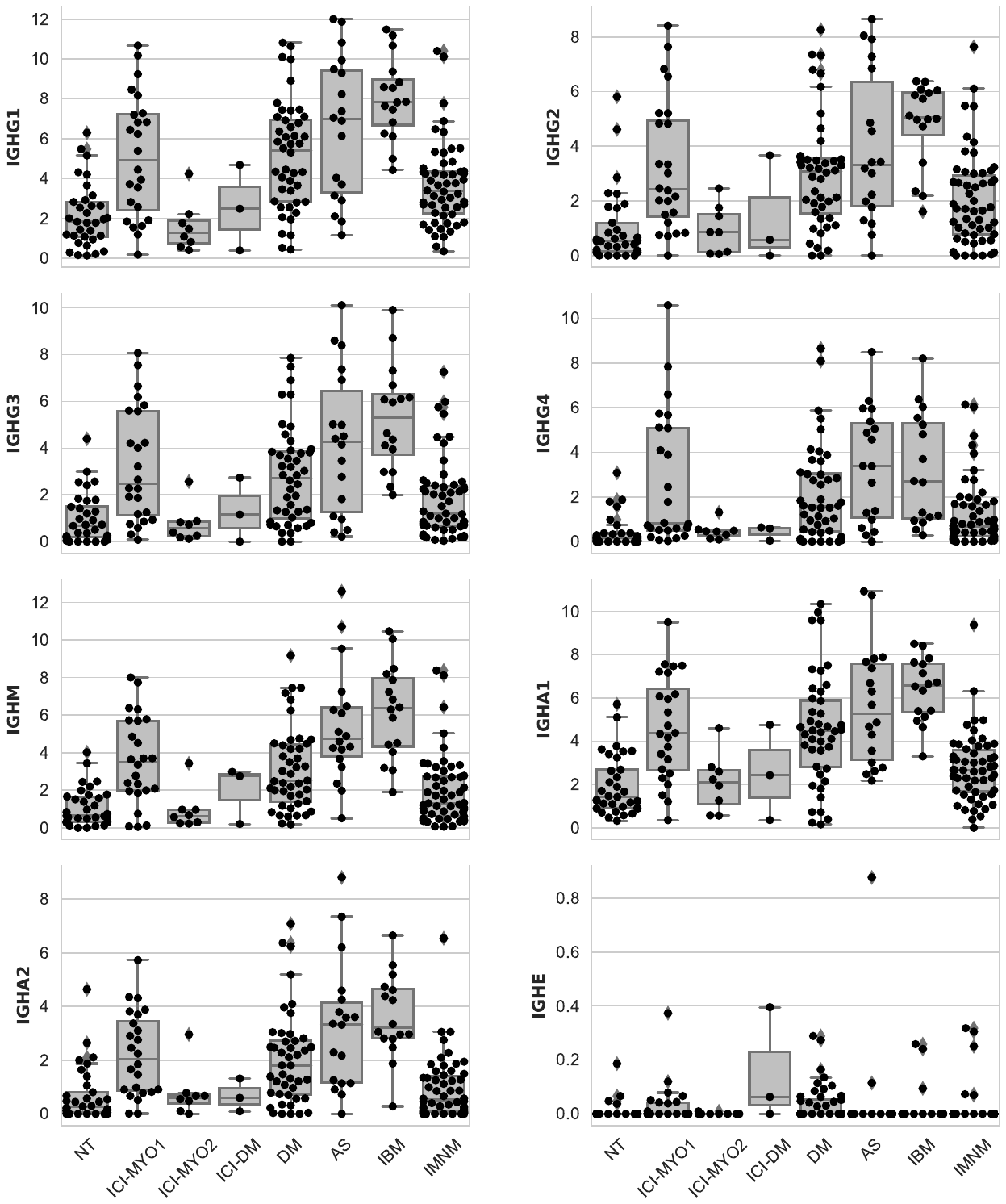

NT: normal muscle; DM: dermatomyositis; AS: antisynthetase syndrome; IBM: inclusion body myositis; IMNM: immune-mediated necrotizing myopathy.

**Supplementary Figure 18.** Expression (average z-score of log2[TMM+1]) of TNF receptors and their ligands showing general overexpression shared amongst different types of inflammatory myopathy.

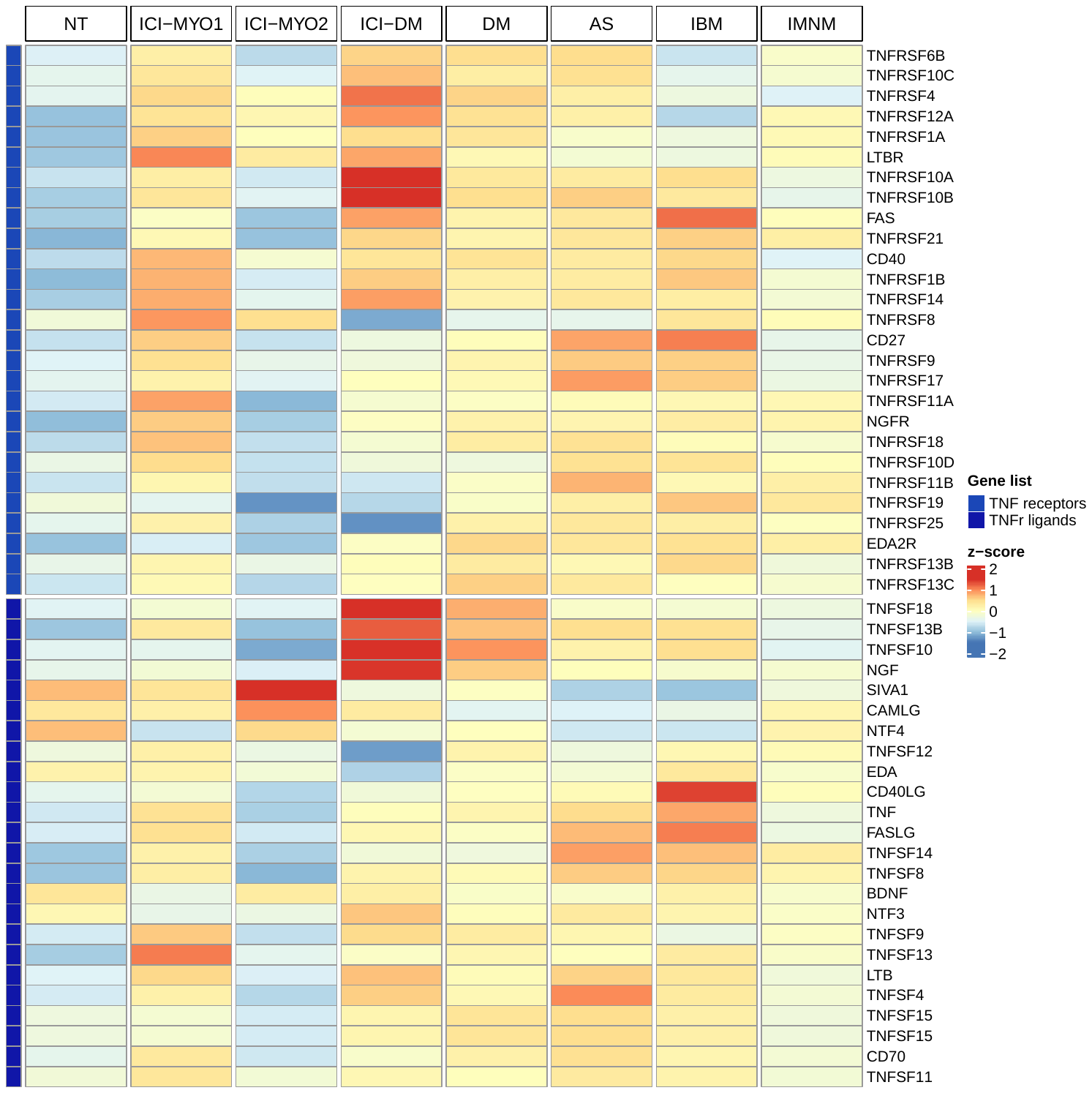

**Figure 19.** Expression (log2[TMM+1]) of representative genes from the TNF pathway.

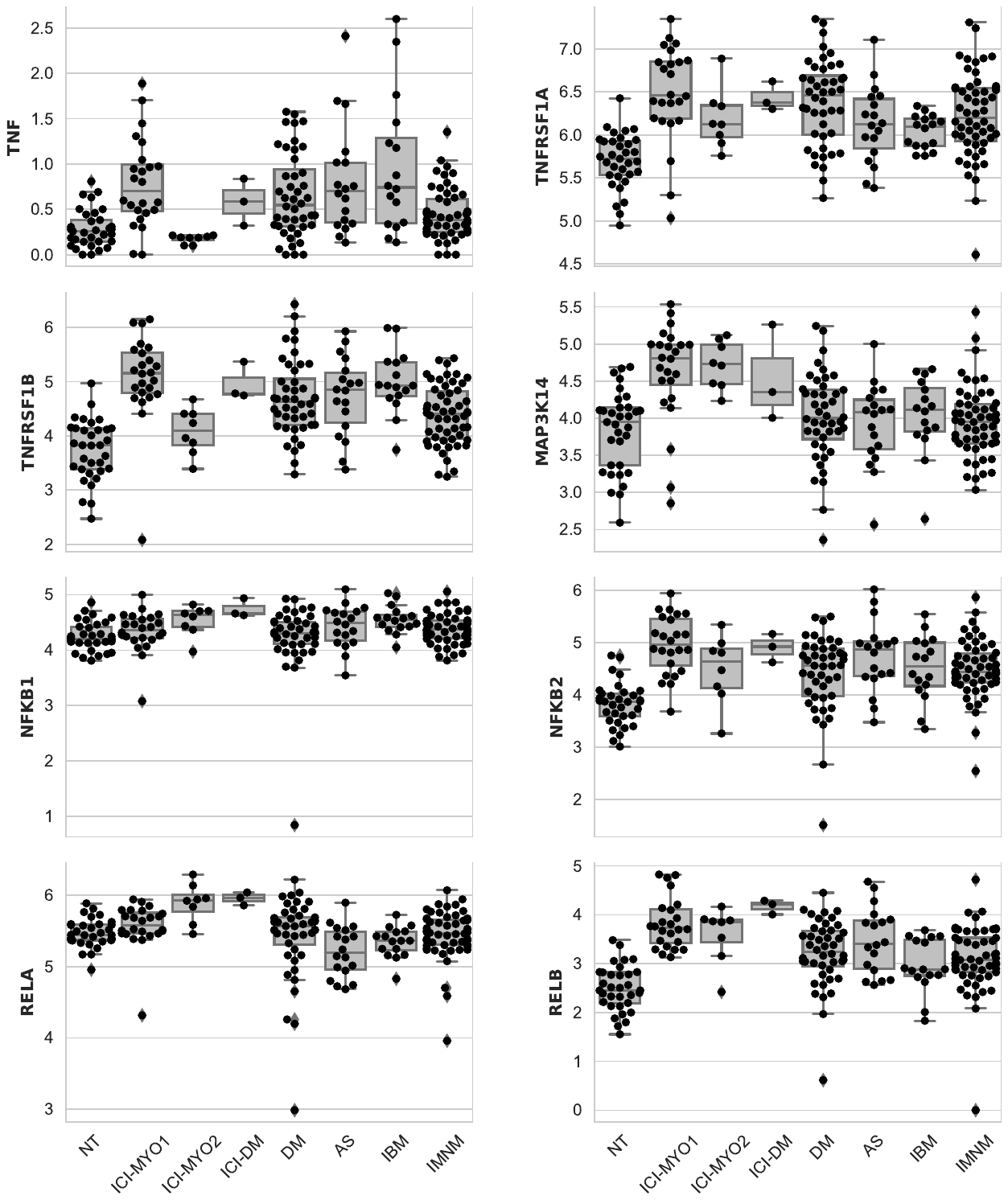

NT: normal muscle; ICI-PM: immune checkpoint-induced myopathy with no skin involvement; ICI-DM: immune checkpoint-induced dermatomyositis; DM: dermatomyositis; AS: antisynthetase syndrome; IBM: inclusion body myositis; IMNM: immune-mediated necrotizing myopathy.

**Supplementary Figure 20.** Gene Set Enrichment Analysis of the TNFA pathway (left) in immune checkpoint-induced myopathy patients compared to normal muscle (p-value 0.01). Fifty genes with the highest signal-to-noise ratio in this pathway (red high, blue low).

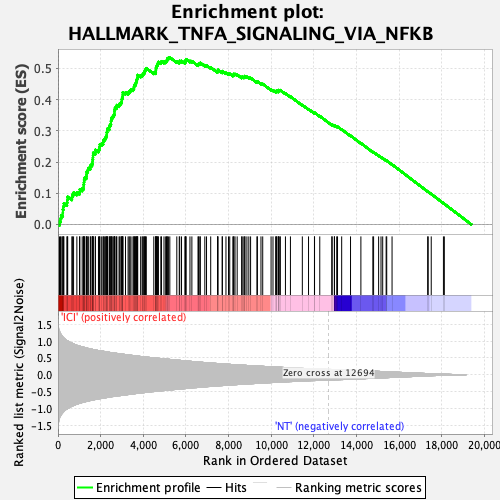

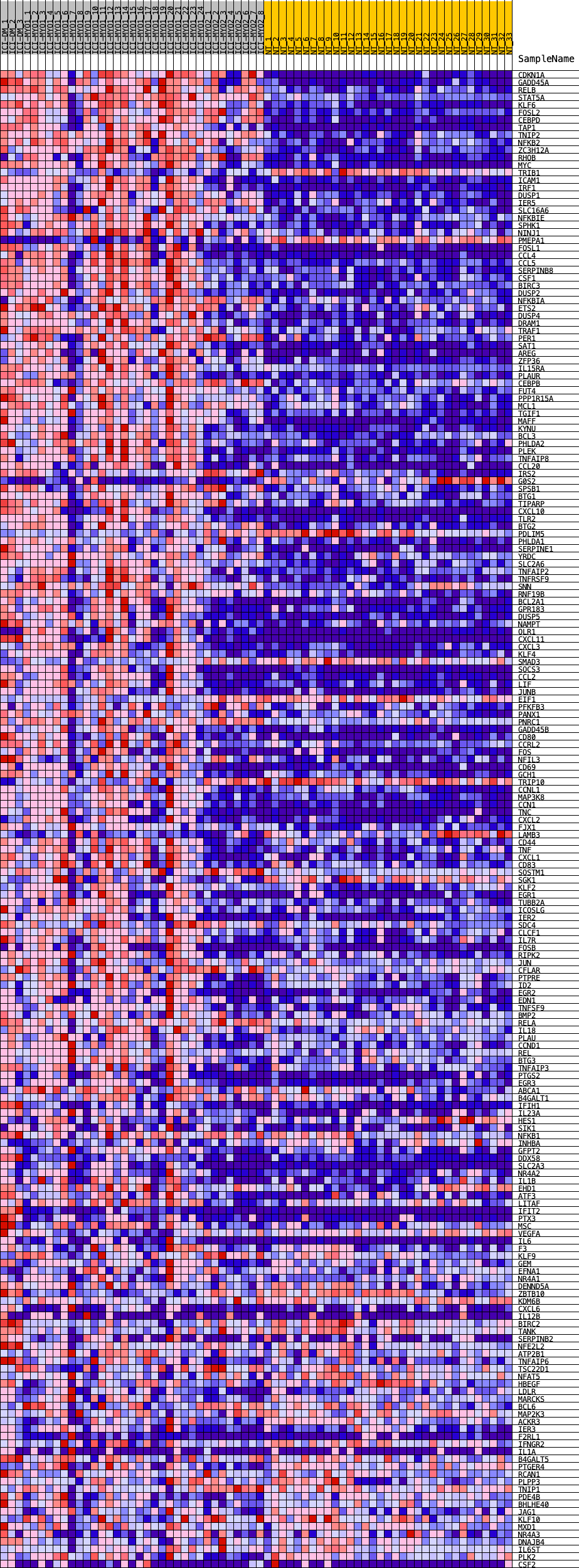

**Supplementary Figure 21.** Expression (log2[TMM+1]) of ICAM1 and VCAM1.

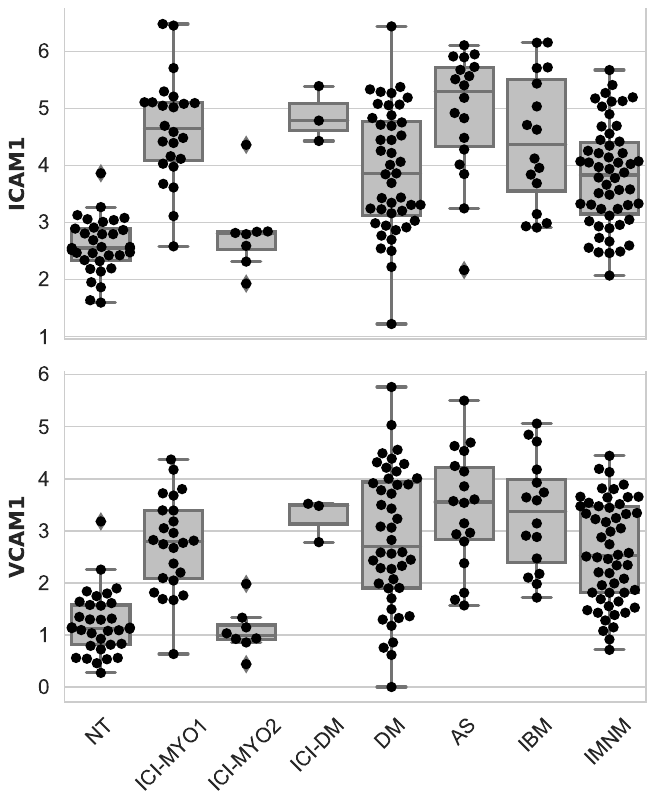

NT: normal muscle; DM: dermatomyositis; AS: antisynthetase syndrome; IBM: inclusion body myositis; IMNM: immune-mediated necrotizing myopathy

**Supplementary Figure 22.** Expression (log2[TMM+1]) of checkpoint genes.

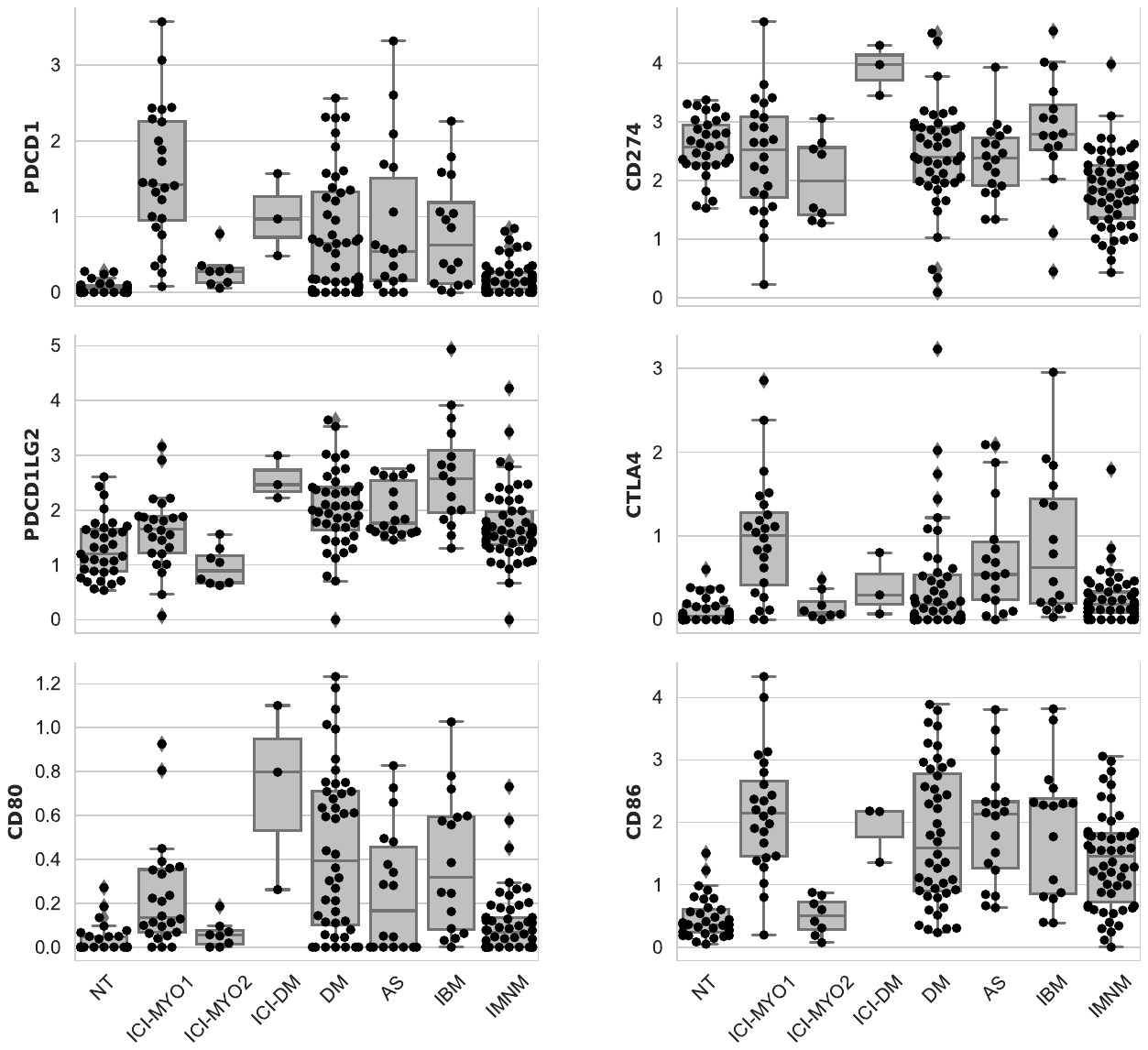

NT: normal muscle; DM: dermatomyositis; AS: antisynthetase syndrome; IBM: inclusion body myositis; IMNM: immune-mediated necrotizing myopathy

**Supplementary Figure 23.** Expression levels (log2[TMM + 1]) of representative genes of the IFNG (row 1) and IL6 (row 2) pathways. Tumors treated with immune checkpoint inhibitors (ICI) have overexpression of IFNG and IFNG-stimulated genes (row 1) but not of genes related to the IL6 pathway (row 2).

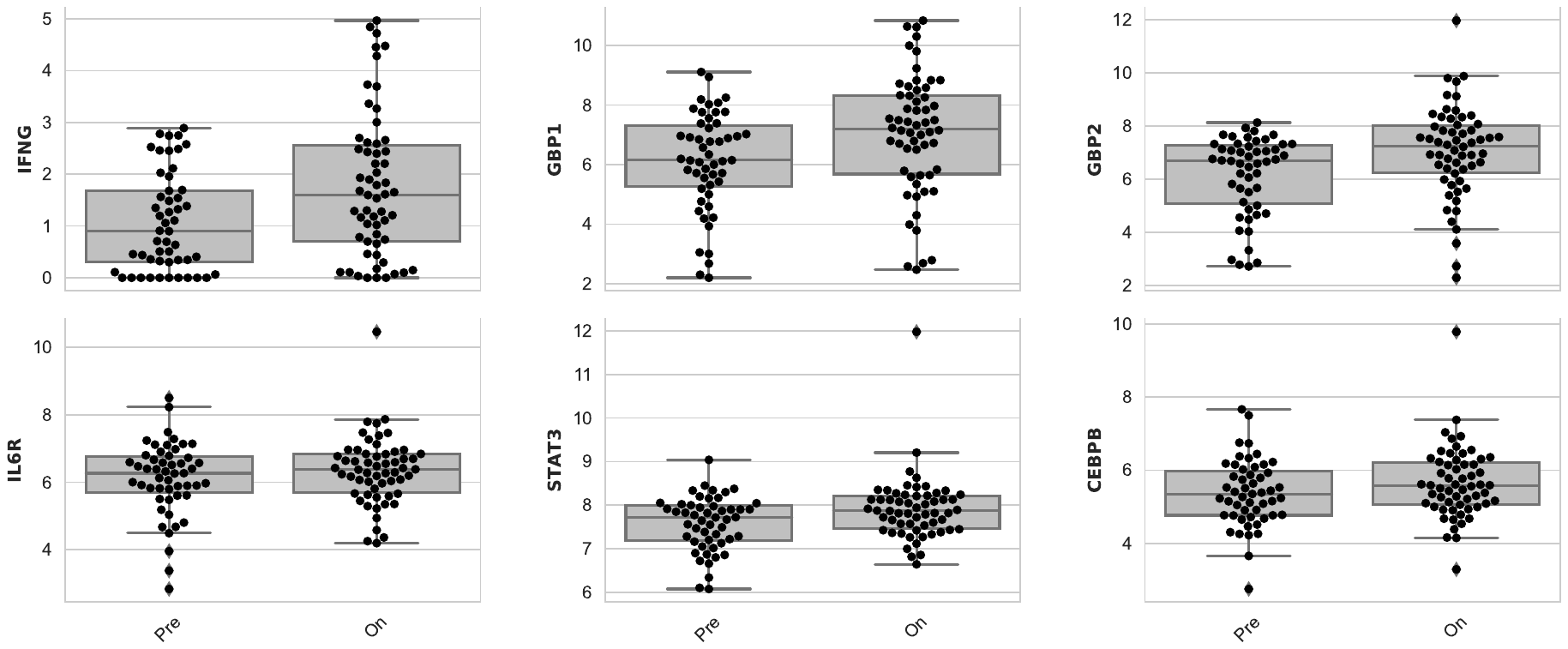

Pre: pre-treatment; On: on-treatment.
