## Supplementary Tables for "Transcriptomic profiling reveals distinct subsets of immune checkpoint inhibitor-induced myositis"

**Supplementary Table 1.** Patient characteristics.

| **Group** | **Female** | **Age at biopsy** | **Dermatomyositis** | **Infl. infiltrates** | **Type of tumor** | **Treat. cycle** | **PD1 inhibitor** | **PD-L1 inhibitor** | **CTLA4 inhibitor** | **Myocarditis** | **Diplopia** | **Dysphagia** | **Anti-striational** | **Anti-AChR** | **Other atbs** | **CK at biopsy** | **Peak CK** |
| --- | --- | --- | --- | --- | --- | --- | --- | --- | --- | --- | --- | --- | --- | --- | --- | --- | --- |
| ICI_DM | no | 73 | yes | yes | Urothelial carcinoma | 2 | yes | no | no | no | no | no | NA | NA | Anti-TIF1g | 428 | 428 |
| ICI_DM | no | 44 | yes | yes | Melanoma | 1 | yes | no | no | no | no | no | no | NA | Anti-TIF1g | 9088 | 9088 |
| ICI_DM | yes | 75 | yes | yes | Urothelial carcinoma | 2 | no | yes | no | no | no | no | no | NA | Anti-TIF1g | 129 | 129 |
| ICI_MYO1 | no | 63 | no | yes | Pancreatic adenocarcinoma | 1 | no | yes | no | yes | yes | no | NA | yes | Anti-Ro60, anti-Ro52, anti-SRP, anti-Mi2a | 2889 | 4204 |
| ICI_MYO1 | no | 69 | no | yes | Liposarcoma | 1 | yes | no | no | no | no | no | no | yes | Anti-MuSK | 172 | 14000 |
| ICI_MYO1 | no | 72 | no | yes | Thyroid anaplastic | 1 | no | yes | yes | yes | yes | no | yes | no | Anti-Jo1 | 912 | 23495 |
| ICI_MYO1 | no | 65 | no | yes | Pancreatic neuroendocrine | 2 | no | yes | yes | no | yes | no | yes | yes | Anti-SRP | 273 | 2430 |
| ICI_MYO1 | no | 82 | no | yes | Melanoma | 1 | yes | no | no | yes | no | no | yes | yes | NA | 85 | 85 |
| ICI_MYO1 | no | 60 | no | yes | Squamous cell lung carcinoma | 3 | no | yes | no | no | no | no | yes | yes | None | 201 | 201 |
| ICI_MYO1 | no | 74 | no | yes | Thymoma | 4 | no | yes | no | no | no | no | yes | no | Anti-PM/Scl100 | 597 | 1150 |
| ICI_MYO1 | no | 63 | no | yes | Squamous cell lung carcinoma | 3 | yes | no | no | no | yes | no | no | no | Anti-TIF1g, anti-Ro52 | 380 | 380 |
| ICI_MYO1 | yes | 61 | no | yes | Squamous head neck carcinoma | 3 | yes | no | no | no | no | no | no | no | Anti-HMGCR | 1900 | 5976 |
| ICI_MYO1 | no | 86 | no | yes | Cholangiocarcinoma | 2 | no | yes | no | yes | no | no | yes | yes | Anti-TIF1g | 638 | 11953 |
| ICI_MYO1 | no | 67 | no | yes | Lung adenocarcinoma | 2 | yes | no | no | yes | no | yes | yes | no | Anti-PM/Scl, anti-M2 | 10850 | 10850 |
| ICI_MYO1 | yes | 62 | no | yes | Breast adenocarcinoma | 1 | yes | no | no | yes | yes | no | yes | yes | None | 11830 | 11830 |
| ICI_MYO1 | yes | 60 | no | yes | Melanoma | 1 | yes | no | no | no | no | no | no | NA | Anti-Mi2 | 1184 | 1184 |
| ICI_MYO1 | yes | 66 | no | yes | Lung adenocarcinoma | 2 | yes | no | no | yes | no | no | yes | yes | Anti-NPX2, anti-Ro52 | 1934 | 1934 |
| ICI_MYO1 | yes | 58 | no | yes | Melanoma | 3 | yes | no | no | no | no | no | NA | NA | None | 666 | 666 |
| ICI_MYO1 | yes | 81 | no | yes | Melanoma | 1 | yes | no | no | yes | no | no | no | no | Anti-NT5c1A | 8333 | 8333 |
| ICI_MYO1 | no | 55 | no | no | Melanoma | 2 | yes | no | no | yes | no | no | no | no | NA | 280 | 1284 |
| ICI_MYO1 | no | 57 | no | yes | Melanoma | 2 | yes | no | no | no | no | no | yes | no | NA | 180 | 444 |
| ICI_MYO1 | no | 49 | no | yes | Esophageal adenocarcinoma | 2 | yes | no | no | yes | yes | no | no | no | NA | 180 | 333 |
| ICI_MYO1 | no | 84 | no | no | Merkel cell carcinoma | 1 | yes | no | no | no | no | yes | yes | yes | None | 1226 | 2685 |
| ICI_MYO1 | no | 78 | no | no | Thymoma | 1 | no | yes | no | yes | no | no | NA | yes | None | 1195 | 1195 |
| ICI_MYO1 | yes | 68 | no | yes | Thymoma | 1 | no | yes | no | yes | no | no | NA | yes | None | 977 | 977 |
| ICI_MYO1 | yes | 59 | no | NA | Thymoma | 2 | no | yes | no | no | no | no | NA | no | None | 1086 | 1086 |
| ICI_MYO1 | no | 50 | no | NA | Thymoma | 1 | no | yes | no | yes | no | no | NA | yes | None | 880 | 1084 |
| ICI_MYO2 | no | 76 | no | yes | Melanoma | 2 | yes | no | no | no | no | no | NA | yes | Anti-Mi2b, anti-NXP2 | 1169 | 4275 |
| ICI_MYO2 | yes | 65 | no | no | Mesothelioma | 3 | yes | no | no | no | no | no | NA | NA | NA | 51 | 481 |
| ICI_MYO2 | no | 69 | no | no | Pancreatic adenocarcinoma | 1 | yes | no | yes | no | no | no | no | no | NA | 5599 | 6260 |
| ICI_MYO2 | no | 69 | no | no | Melanoma | 1 | yes | no | no | no | no | no | yes | no | NA | 427 | 7307 |
| ICI_MYO2 | no | 46 | no | no | Follicular lymphoma | 2 | yes | no | no | no | yes | no | yes | yes | NA | 72 | 72 |
| ICI_MYO2 | no | 70 | no | yes | Melanoma | 1 | yes | no | no | no | no | yes | yes | no | None | NA | 542 |
| ICI_MYO2 | yes | 88 | no | no | Renal cell carcinoma | 2 | yes | no | yes | no | no | yes | no | no | None | 26 | 28 |
| ICI_MYO2 | no | 68 | no | yes | Renal cell carcinoma | 1 | yes | no | yes | no | no | no | yes | no | NA | 492 | 762 |

**Supplementary Table 2.** Type of immune checkpoint inhibitor.

| **Group** | **Nivolumab** | **Pembrolizumab** | **Atezolizumab** | **Avelumab** | **Durvalumab** | **Tremelimumab** | **Ipilimumab** | **M7824** |
| --- | --- | --- | --- | --- | --- | --- | --- | --- |
| ICI_DM | no | yes | no | no | no | no | no | no |
| ICI_DM | yes | no | no | no | no | no | no | no |
| ICI_DM | no | no | no | yes | no | no | no | no |
| ICI_MYO1 | no | no | yes | no | no | no | no | no |
| ICI_MYO1 | no | yes | no | no | no | no | no | no |
| ICI_MYO1 | no | no | no | no | yes | yes | no | no |
| ICI_MYO1 | no | no | no | no | yes | yes | no | no |
| ICI_MYO1 | no | yes | no | no | no | no | no | no |
| ICI_MYO1 | no | no | no | no | yes | no | no | no |
| ICI_MYO1 | no | no | yes | no | no | no | no | no |
| ICI_MYO1 | yes | no | no | no | no | no | no | no |
| ICI_MYO1 | no | yes | no | no | no | no | no | no |
| ICI_MYO1 | no | no | no | no | no | no | no | yes |
| ICI_MYO1 | no | yes | no | no | no | no | no | no |
| ICI_MYO1 | no | yes | no | no | no | no | no | no |
| ICI_MYO1 | yes | no | no | no | no | no | no | no |
| ICI_MYO1 | no | yes | no | no | no | no | no | no |
| ICI_MYO1 | yes | no | no | no | no | no | no | no |
| ICI_MYO1 | yes | no | no | no | no | no | no | no |
| ICI_MYO1 | no | yes | no | no | no | no | no | no |
| ICI_MYO1 | no | yes | no | no | no | no | no | no |
| ICI_MYO1 | no | yes | no | no | no | no | no | no |
| ICI_MYO1 | no | yes | no | no | no | no | no | no |
| ICI_MYO1 | no | no | no | yes | no | no | no | no |
| ICI_MYO1 | no | no | no | yes | no | no | no | no |
| ICI_MYO1 | no | no | no | yes | no | no | no | no |
| ICI_MYO1 | no | no | no | yes | no | no | no | no |
| ICI_MYO2 | no | yes | no | no | no | no | no | no |
| ICI_MYO2 | no | yes | no | no | no | no | no | no |
| ICI_MYO2 | yes | no | no | no | no | no | yes | no |
| ICI_MYO2 | no | yes | no | no | no | no | no | no |
| ICI_MYO2 | no | yes | no | no | no | no | no | no |
| ICI_MYO2 | yes | no | no | no | no | no | no | no |
| ICI_MYO2 | yes | no | no | no | no | no | yes | no |
| ICI_MYO2 | yes | no | no | no | no | no | yes | no |

**Supplementary Table 3.** Differential expression of relevant genes related to immune-checkpoint inhibitor myopathy by bulk RNA sequencing. The different groups identified by the unsupervised clustering analysis (ICI-DM, ICI-MYO1, and ICI-MYO2) were compared to normal muscle biopsies (NT) and to each other. Missing values correspond to genes that did not pass the cutoff to be included in the differential expression.

| **Gene** | **ICI-MYO1**  **vs.**  **NT** | | **ICI-MYO2**  **vs.**  **NT** | | **ICI-DM**  **vs.**  **NT** | | **ICI-MYO1**  **vs.**  **ICI-MYO2** | | **ICI-DM**  **vs.**  **ICI-MYO1** | | **ICI-DM**  **vs.**  **ICI-MYO2** | |
| --- | --- | --- | --- | --- | --- | --- | --- | --- | --- | --- | --- | --- |
|  | **logFC** | **adj.P.Val** | **logFC** | **adj.P.Val** | **logFC** | **adj.P.Val** | **logFC** | **adj.P.Val** | **logFC** | **adj.P.Val** | **logFC** | **adj.P.Val** |
| ACTA1 | -2.2 | 5e-08 | -0.8 | 2e-05 | -3.4 | 7e-16 | -1.3 | 0.1 | -1.2 | 0.6 | -2.6 | 0.002 |
| BDNF | -1.1 | 0.07 | 0.1 | 0.9 | 0.3 | 0.8 | -1.3 | 0.07 | 1.5 | 0.3 | 0.1 | 0.9 |
| CAMLG | -0.0 | 0.6 | 0.2 | 0.1 | -0.0 | 1 | -0.2 | 0.1 | 0.1 | 0.9 | -0.2 | 0.3 |
| CD14 | 2.3 | 1e-07 | -0.7 | 0.05 | 1.9 | 2e-05 | 3.0 | 0.002 | -0.5 | 0.8 | 2.5 | 9e-04 |
| CD19 |  | NA |  | NA |  | NA | 2.5 | 0.007 | -0.0 | 1 |  | NA |
| CD27 | 2.8 | 1e-05 |  | NA | 1.4 | 0.03 | 3.2 | 0.008 | -1.5 | 0.6 | 1.7 | 0.2 |
| CD274 | -0.3 | 0.4 | -0.9 | 0.004 | 1.5 | 6e-06 | 0.5 | 0.6 | 1.9 | 0.1 | 2.3 | 0.003 |
| CD3E | 1.9 | 0.01 | -0.4 | 0.5 | 1.8 | 0.001 | 2.2 | 0.2 | -0.2 | 1 | 2.0 | 0.1 |
| CD4 | 2.8 | 2e-11 | -0.7 | 0.06 | 1.5 | 8e-04 | 3.5 | 2e-04 | -1.2 | 0.3 | 2.2 | 0.01 |
| CD40 | 0.9 | 8e-08 | 0.3 | 0.03 | 0.7 | 4e-04 | 0.6 | 0.04 | -0.2 | 0.8 | 0.4 | 0.2 |
| CD40LG |  | NA |  | NA | 0.4 | 0.7 | 2.7 | 0.02 | 0.0 | 1 |  | NA |
| CD68 | 3.1 | 1e-08 | 0.5 | 0.2 | 3.7 | 1e-09 | 2.5 | 0.01 | 0.6 | 0.8 | 3.1 | 0.001 |
| CD70 |  | NA |  | NA | 1.5 | 0.06 | 3.0 | 0.002 | -0.5 | 0.8 |  | NA |
| CD80 |  | NA |  | NA |  | NA | 2.2 | 0.02 | 2.2 | 0.04 | 4.4 | 0.002 |
| CD86 | 3.1 | 1e-06 | -0.2 | 0.7 | 3.1 | 5e-10 | 3.2 | 0.007 | 0.1 | 1 | 3.2 | 0.002 |
| CD8A | 3.5 | 8e-06 | 0.5 | 0.3 | 2.4 | 6e-05 | 3.0 | 0.03 | -1.2 | 0.7 | 1.7 | 0.2 |
| CEBPB | 0.8 | 5e-04 | 1.4 | 1e-04 | 1.0 | 0.05 | -0.6 | 0.2 | 0.2 | 0.9 | -0.5 | 0.4 |
| COX7B | -1.2 | 1e-14 | -0.7 | 2e-04 | -1.7 | 7e-09 | -0.6 | 0.04 | -0.4 | 0.5 | -1.1 | 0.02 |
| COX7C | -0.9 | 1e-12 | -0.3 | 0.05 | -1.2 | 7e-06 | -0.6 | 0.006 | -0.3 | 0.6 | -0.9 | 0.006 |
| CTLA4 |  | NA |  | NA |  | NA | 3.2 | 0.04 | -1.2 | 0.7 | 2.0 | 0.2 |
| EDA | 0.2 | 0.5 | -0.4 | 0.05 | -0.7 | 0.007 | 0.5 | 0.3 | -0.8 | 0.4 | -0.4 | 0.5 |
| EDA2R | 0.9 | 0.005 | 0.1 | 0.9 | 1.5 | 0.001 | 0.8 | 0.2 | 0.7 | 0.5 | 1.5 | 0.1 |
| EGR1 | 2.8 | 2e-05 | 0.1 | 0.9 | 3.2 | 0.01 | 2.7 | 0.01 | 0.2 | 1 | 3.0 | 0.03 |
| FAS | 0.8 | 2e-04 | -0.2 | 0.5 | 1.5 | 2e-06 | 0.9 | 0.02 | 0.8 | 0.2 | 1.7 | 0.003 |
| FASLG |  | NA |  | NA | 2.7 | 6e-05 | 2.5 | 0.05 | 0.1 | 1 | 2.7 | 0.006 |
| FOS | 3.5 | 9e-06 | 0.7 | 0.5 | 3.6 | 0.02 | 2.9 | 0.01 | -0.0 | 1 | 2.8 | 0.05 |
| FOSB | 4.6 | 4e-06 | 1.0 | 0.2 | 3.5 | 5e-04 | 3.8 | 0.03 | -1.3 | 0.7 | 2.6 | 0.05 |
| FOSL1 | 4.2 | 2e-08 | 1.4 | 0.006 | 5.8 | 7e-15 | 2.7 | 0.005 | 1.7 | 0.1 | 4.3 | 4e-07 |
| GBP2 | 2.5 | 1e-13 | 1.2 | 6e-06 | 3.0 | 1e-11 | 1.3 | 0.04 | 0.5 | 0.8 | 1.8 | 0.01 |
| GZMA | 2.3 | 0.004 | -0.3 | 0.5 | 1.9 | 4e-04 | 2.4 | 0.1 | -0.5 | 0.9 | 2.0 | 0.08 |
| GZMB | 2.7 | 0.009 | 0.4 | 0.5 | 3.7 | 3e-09 | 2.2 | 0.2 | 1.0 | 0.8 | 3.1 | 0.004 |
| ICAM1 | 2.3 | 4e-13 | 0.1 | 0.8 | 2.5 | 1e-10 | 2.2 | 2e-04 | 0.2 | 0.9 | 2.4 | 7e-04 |
| IFI30 | 5.0 | 4e-13 | 0.4 | 0.7 | 5.2 | 2e-10 | 4.6 | 9e-04 | 0.2 | 0.9 | 4.7 | 0.02 |
| IFNA1 |  | NA |  | NA |  | NA |  | NA |  | NA |  | NA |
| IFNA10 |  | NA |  | NA |  | NA |  | NA |  | NA |  | NA |
| IFNA13 |  | NA |  | NA |  | NA |  | NA |  | NA |  | NA |
| IFNA14 |  | NA |  | NA |  | NA |  | NA |  | NA |  | NA |
| IFNA16 |  | NA |  | NA |  | NA |  | NA |  | NA |  | NA |
| IFNA17 |  | NA |  | NA |  | NA |  | NA |  | NA |  | NA |
| IFNA2 |  | NA |  | NA |  | NA |  | NA |  | NA |  | NA |
| IFNA21 |  | NA |  | NA |  | NA |  | NA |  | NA |  | NA |
| IFNA4 |  | NA |  | NA |  | NA |  | NA |  | NA |  | NA |
| IFNA5 |  | NA |  | NA |  | NA |  | NA |  | NA |  | NA |
| IFNA6 |  | NA |  | NA |  | NA |  | NA |  | NA |  | NA |
| IFNA7 |  | NA |  | NA |  | NA |  | NA |  | NA |  | NA |
| IFNA8 |  | NA |  | NA |  | NA |  | NA |  | NA |  | NA |
| IFNAR1 | -0.5 | 7e-06 | -0.4 | 0.003 | -0.4 | 0.1 | -0.1 | 0.7 | 0.2 | 0.7 | 0.0 | 0.9 |
| IFNAR2 | 0.4 | 4e-04 | -0.5 | 0.002 | -1.5 | 0.5 | 0.9 | 8e-04 | -1.4 | 0.3 | 0.5 | 0.09 |
| IFNB1 |  | NA |  | NA |  | NA |  | NA |  | NA |  | NA |
| IFNG |  | NA |  | NA |  | NA | 3.5 | 0.03 | -1.7 | 0.7 | 1.9 | 0.3 |
| IGHA1 | 3.0 | 2e-04 | -0.3 | 0.7 | 0.9 | 0.5 | 3.2 | 0.05 | -2.1 | 0.6 | 1.0 | 0.6 |
| IGHA2 | 2.5 | 0.05 | 0.1 | 1 | 1.2 | 0.4 | 2.3 | 0.3 | -1.5 | 0.8 | 0.9 | 0.7 |
| IGHE |  | NA |  | NA |  | NA |  | NA |  | NA |  | NA |
| IGHG1 | 3.4 | 6e-04 | -1.2 | 0.2 | 0.8 | 0.6 | 4.4 | 0.04 | -2.6 | 0.6 | 1.7 | 0.3 |
| IGHG2 | 3.0 | 0.03 | -0.6 | 0.6 | 0.8 | 0.7 | 3.2 | 0.2 | -2.6 | 0.8 | 0.8 | 0.8 |
| IGHG3 | 3.3 | 0.008 | -0.3 | 0.8 | 0.9 | 0.6 | 3.4 | 0.1 | -2.8 | 0.7 | 0.8 | 0.7 |
| IGHG4 | 3.9 | 0.02 | 0.7 | 0.3 | 1.1 | 0.4 | 2.9 | 0.3 | -2.9 | 0.7 | 0.1 | 0.9 |
| IGHM | 2.9 | 0.01 | -0.8 | 0.3 | 1.7 | 0.08 | 3.4 | 0.1 | -1.1 | 0.9 | 2.2 | 0.2 |
| IL17RC | 0.5 | 0.01 | 0.8 | 0.02 | -0.1 | 0.9 | -0.3 | 0.2 | -0.6 | 0.3 | -0.9 | 0.05 |
| IL6 | 4.4 | 4e-05 |  | NA | 5.2 | 1e-10 | 3.4 | 0.04 | 0.8 | 0.8 | 4.2 | 0.002 |
| IL6R | 1.3 | 4e-07 | 1.9 | 2e-07 | 0.9 | 0.02 | -0.7 | 0.2 | -0.3 | 0.8 | -1.1 | 0.2 |
| ISG15 | 2.1 | 5e-09 | 0.4 | 0.2 | 9.0 | 3e-22 | 1.6 | 0.006 | 6.9 | 2e-05 | 8.5 | 1e-08 |
| JUN | 0.5 | 0.04 | 0.4 | 0.1 | 0.7 | 0.07 | 0.1 | 0.8 | 0.2 | 0.9 | 0.3 | 0.5 |
| JUNB | 3.1 | 1e-08 | 0.7 | 0.3 | 3.1 | 8e-04 | 2.5 | 0.007 | -0.1 | 1 | 2.3 | 0.01 |
| LTB | 2.4 | 4e-06 | 0.2 | 0.7 | 2.9 | 6e-05 | 2.1 | 0.009 | 0.5 | 0.8 | 2.5 | 0.01 |
| LTBR | 1.2 | 5e-10 | 0.7 | 4e-05 | 1.1 | 3e-05 | 0.5 | 0.2 | -0.1 | 0.9 | 0.3 | 0.2 |
| MAP3K14 | 0.9 | 1e-05 | 1.0 | 2e-04 | 0.8 | 0.04 | -0.1 | 0.8 | -0.0 | 1 | -0.2 | 0.6 |
| MS4A1 |  | NA |  | NA | 2.1 | 0.02 | 2.6 | 0.07 | 0.9 | 0.8 | 3.5 | 0.04 |
| MT-TI | -2.2 | 1e-08 | -1.5 | 0.001 | -2.7 | 0.005 | -0.7 | 0.1 | -0.4 | 0.8 | -1.1 | 0.1 |
| MT-TL1 | -2.9 | 5e-17 | -2.2 | 1e-07 | -2.8 | 6e-06 | -0.8 | 0.04 | 0.2 | 0.9 | -0.6 | 0.3 |
| MX1 | 0.7 | 0.007 | -0.6 | 0.03 | 7.0 | 2e-19 | 1.3 | 0.02 | 6.3 | 2e-05 | 7.6 | 4e-08 |
| MYH1 | -1.0 | 0.09 | 0.3 | 0.7 | -2.9 | 0.008 | -1.3 | 0.3 | -1.8 | 0.4 | -3.2 | 0.02 |
| MYH3 | 2.7 | 5e-07 | 1.0 | 0.07 | 0.8 | 0.3 | 1.8 | 0.07 | -1.9 | 0.3 | -0.3 | 0.8 |
| NCAM1 | 1.5 | 3e-06 | -0.0 | 0.9 | 1.1 | 0.02 | 1.5 | 0.01 | -0.4 | 0.8 | 1.1 | 0.05 |
| NFKB1 | 0.1 | 0.3 | 0.3 | 0.03 | 0.5 | 0.003 | -0.2 | 0.4 | 0.5 | 0.3 | 0.2 | 0.4 |
| NFKB2 | 1.2 | 4e-11 | 0.7 | 0.004 | 1.2 | 1e-04 | 0.5 | 0.09 | -0.0 | 1 | 0.4 | 0.4 |
| NGF | 0.1 | 0.7 | -0.4 | 0.4 | 1.9 | 2e-05 | 0.5 | 0.3 | 1.8 | 0.02 | 2.3 | 0.004 |
| NGFR | 2.9 | 2e-09 | 0.2 | 0.8 | 2.0 | 0.004 | 2.7 | 8e-04 | -0.9 | 0.5 | 1.7 | 0.09 |
| NTF3 | -0.4 | 0.1 | -0.8 | 0.04 | 0.3 | 0.6 | 0.2 | 0.6 | 0.8 | 0.4 | 1.0 | 0.3 |
| NTF4 | -1.4 | 4e-05 | -0.2 | 0.4 | -0.7 | 0.03 | -1.2 | 0.06 | 0.8 | 0.7 | -0.5 | 0.4 |
| PDCD1 | 5.1 | 4e-10 |  | NA | 4.6 | 2e-12 | 3.1 | 0.006 | -0.6 | 0.8 | 2.5 | 0.01 |
| PDCD1LG2 | 0.3 | 0.6 | -0.9 | 0.01 | 1.7 | 1e-05 | 1.0 | 0.2 | 1.5 | 0.3 | 2.5 | 4e-04 |
| PRF1 | 2.1 | 2e-04 | 0.4 | 0.3 | 2.8 | 1e-06 | 1.6 | 0.1 | 0.7 | 0.8 | 2.3 | 0.008 |
| RELA | 0.1 | 0.1 | 0.4 | 7e-04 | 0.5 | 0.001 | -0.3 | 0.1 | 0.4 | 0.2 | 0.1 | 0.8 |
| RELB | 1.5 | 2e-13 | 1.3 | 2e-06 | 1.9 | 1e-08 | 0.3 | 0.4 | 0.4 | 0.5 | 0.6 | 0.2 |
| SIVA1 | -0.2 | 0.2 | 0.5 | 0.02 | -0.6 | 0.05 | -0.7 | 0.02 | -0.4 | 0.4 | -1.1 | 0.01 |
| SOCS3 | 4.4 | 2e-11 | 1.2 | 0.02 | 4.5 | 4e-08 | 3.3 | 0.001 | -0.0 | 1 | 3.3 | 0.002 |
| STAT3 | 0.4 | 4e-05 | 0.5 | 5e-04 | 0.8 | 5e-04 | -0.2 | 0.4 | 0.4 | 0.3 | 0.2 | 0.5 |
| TGFB1 | 1.6 | 2e-10 | -0.1 | 0.6 | 0.8 | 0.01 | 1.7 | 4e-04 | -0.7 | 0.4 | 1.0 | 0.03 |
| TGFB2 | -0.5 | 0.1 | -1.0 | 0.002 | 1.2 | 0.002 | 0.5 | 0.4 | 1.7 | 0.06 | 2.2 | 0.005 |
| TGFB3 | 0.1 | 0.8 | -0.0 | 1 | 1.1 | 1e-04 | 0.0 | 1 | 1.1 | 0.3 | 1.1 | 0.02 |
| TGFBR2 | 0.9 | 1e-04 | 0.1 | 0.7 | 0.8 | 0.06 | 0.8 | 0.07 | -0.1 | 0.9 | 0.6 | 0.06 |
| TNF | 1.5 | 0.01 | -0.5 | 0.3 | 1.6 | 0.008 | 2.0 | 0.07 | 0.0 | 1 | 2.0 | 0.004 |
| TNFRSF10A | 1.3 | 1e-04 | 0.2 | 0.6 | 2.7 | 2e-10 | 1.1 | 0.04 | 1.4 | 0.07 | 2.5 | 4e-05 |
| TNFRSF10B | 1.0 | 2e-07 | 0.3 | 0.2 | 1.9 | 1e-09 | 0.7 | 0.03 | 0.9 | 0.08 | 1.6 | 0.002 |
| TNFRSF10C | 2.0 | 3e-05 |  | NA | 2.6 | 2e-04 | 1.6 | 0.02 | 0.5 | 0.7 | 2.2 | 0.02 |
| TNFRSF10D | 0.8 | 0.003 | -0.4 | 0.1 | 0.1 | 0.8 | 1.2 | 0.03 | -0.7 | 0.6 | 0.4 | 0.4 |
| TNFRSF11A | 2.0 | 6e-06 | -1.3 | 0.005 | 0.7 | 0.1 | 3.2 | 9e-04 | -1.2 | 0.5 | 2.0 | 0.02 |
| TNFRSF11B | 1.7 | 2e-04 | 0.3 | 0.6 | 0.3 | 0.8 | 1.4 | 0.02 | -1.4 | 0.3 | -0.0 | 1 |
| TNFRSF12A | 2.1 | 2e-08 | 1.7 | 0.001 | 2.9 | 1e-04 | 0.4 | 0.4 | 0.7 | 0.5 | 1.2 | 0.06 |
| TNFRSF13B |  | NA |  | NA |  | NA |  | NA |  | NA |  | NA |
| TNFRSF13C | 1.4 | 2e-08 | -0.3 | 0.4 | 1.1 | 0.006 | 1.8 | 2e-04 | -0.2 | 0.8 | 1.5 | 0.02 |
| TNFRSF14 | 1.5 | 5e-07 | 0.4 | 0.2 | 1.6 | 1e-04 | 1.1 | 0.05 | 0.1 | 1 | 1.2 | 0.08 |
| TNFRSF17 |  | NA |  | NA |  | NA | 1.6 | 0.2 | 0.8 | 0.8 |  | NA |
| TNFRSF18 | 3.1 | 7e-10 |  | NA | 2.2 | 5e-04 | 2.8 | 6e-04 | -1.0 | 0.5 | 1.8 | 0.04 |
| TNFRSF19 | -0.0 | 0.9 | -1.6 | 5e-05 | -0.6 | 0.1 | 1.5 | 0.008 | -0.5 | 0.6 | 0.9 | 0.4 |
| TNFRSF1A | 0.8 | 1e-08 | 0.5 | 0.004 | 0.8 | 0.001 | 0.3 | 0.2 | -0.0 | 1 | 0.3 | 0.4 |
| TNFRSF1B | 1.5 | 1e-09 | 0.3 | 0.2 | 1.3 | 4e-04 | 1.1 | 0.008 | -0.1 | 0.9 | 0.9 | 0.02 |
| TNFRSF21 | 1.1 | 2e-07 | 0.1 | 0.8 | 1.4 | 2e-06 | 1.0 | 0.005 | 0.4 | 0.6 | 1.3 | 0.006 |
| TNFRSF25 | 0.6 | 6e-04 | -0.3 | 0.2 | -0.7 | 0.1 | 0.9 | 0.003 | -1.2 | 0.03 | -0.4 | 0.5 |
| TNFRSF4 | 1.2 | 0.001 | 0.8 | 0.06 | 2.2 | 6e-06 | 0.4 | 0.6 | 1.0 | 0.4 | 1.4 | 0.1 |
| TNFRSF6B | 0.7 | 0.3 | -2.1 | 0.1 | 1.8 | 0.02 | 2.9 | 0.2 | 1.1 | 0.7 | 3.9 | 0.1 |
| TNFRSF8 | 1.2 | 0.001 | 0.7 | 0.03 | -1.5 | 0.01 | 0.4 | 0.5 | -2.6 | 0.1 | -2.2 | 0.05 |
| TNFRSF9 |  | NA |  | NA |  | NA | 1.7 | 0.04 | -1.3 | 0.5 | 0.4 | 0.7 |
| TNFSF10 | -0.0 | 1 | -0.8 | 6e-04 | 1.8 | 6e-08 | 0.8 | 0.2 | 1.9 | 0.1 | 2.6 | 1e-04 |
| TNFSF11 |  | NA |  | NA |  | NA |  | NA |  | NA |  | NA |
| TNFSF12 | 0.3 | 0.07 | -0.1 | 0.7 | -0.5 | 0.06 | 0.3 | 0.3 | -0.8 | 0.3 | -0.5 | 0.3 |
| TNFSF13 | 2.1 | 7e-11 | 0.6 | 0.04 | 1.0 | 0.01 | 1.5 | 0.002 | -1.1 | 0.3 | 0.4 | 0.2 |
| TNFSF13B | 2.3 | 3e-05 | -0.4 | 0.3 | 3.8 | 2e-14 | 2.5 | 0.02 | 1.6 | 0.3 | 4.1 | 2e-05 |
| TNFSF14 | 2.4 | 1e-05 | 0.3 | 0.6 | 2.0 | 0.002 | 2.0 | 0.02 | -0.4 | 0.9 | 1.6 | 0.04 |
| TNFSF15 | 0.2 | 0.4 | -0.7 | 0.1 | 0.7 | 0.3 | 0.9 | 0.03 | 0.5 | 0.6 | 1.3 | 0.1 |
| TNFSF18 |  | NA |  | NA | 5.3 | 1e-15 |  | NA | 4.0 | 2e-05 | 5.7 | 7e-05 |
| TNFSF4 | 1.3 | 2e-05 | -0.3 | 0.5 | 1.8 | 3e-04 | 1.6 | 0.002 | 0.5 | 0.6 | 2.0 | 0.01 |
| TNFSF8 | 1.9 | 0.01 | -0.5 | 0.4 | 2.4 | 1e-05 | 2.3 | 0.08 | 0.5 | 0.9 | 2.8 | 0.002 |
| TNFSF9 | 1.8 | 4e-07 | -0.2 | 0.6 | 1.8 | 2e-04 | 2.0 | 0.001 | -0.0 | 1 | 2.0 | 0.005 |
| TYK2 | 0.3 | 0.01 | 0.5 | 0.002 | -0.1 | 0.8 | -0.2 | 0.3 | -0.3 | 0.6 | -0.6 | 0.04 |
| VCAM1 | 2.2 | 2e-07 | -0.4 | 0.3 | 2.8 | 5e-09 | 2.5 | 0.003 | 0.7 | 0.6 | 3.2 | 1e-04 |

**Supplementary Table 4.** Differential expression of relevant genes related to immune-checkpoint inhibitor myopathy by single-nuclei RNA sequencing. Four biopsies from the cluster ICI-MYO1 were compared with a representative selection of patients with other types of inflammatory myopathy (4 dermatomyositis, 3 antisynthetase syndrome, 6 immune-mediated necrotizing myositis, and 2 inclusion body myositis). Missing values correspond to genes that did not pass the cutoff to be included in the differential expression.

| **Gene** | **T-cells**  **ICI-MYO1 vs.**  **other myositis** | | **FAP cells**  **ICI-MYO1 vs.**  **other myositis** | | **Adipocytes**  **ICI-MYO1 vs.**  **other myositis** | | **Endothelial cells**  **ICI-MYO1 vs.**  **other myositis** | | **Myeloid cells**  **ICI-MYO1 vs.**  **other myositis** | | **Fibroblasts**  **ICI-MYO1 vs.**  **other myositis** | | **Muscle fibers**  **ICI-MYO1 vs.**  **other myositis** | | **Satellite cells**  **ICI-MYO1 vs.**  **other myositis** | |
| --- | --- | --- | --- | --- | --- | --- | --- | --- | --- | --- | --- | --- | --- | --- | --- | --- |
|  | **avg_log2FC** | **p_val_adj** | **avg_log2FC** | **p_val_adj** | **avg_log2FC** | **p_val_adj** | **avg_log2FC** | **p_val_adj** | **avg_log2FC** | **p_val_adj** | **avg_log2FC** | **p_val_adj** | **avg_log2FC** | **p_val_adj** | **avg_log2FC** | **p_val_adj** |
| CEBPB |  | NA |  | NA | 0.3 | 0.1 |  | NA |  | NA |  | NA | 0.3 | 6e-35 | 0.6 | 2e-07 |
| EGR1 | 0.3 | 0.7 | 1.0 | 5e-13 | 0.7 | 7e-32 | 0.6 | 4e-20 | 0.9 | 3e-25 | 1.0 | 2e-94 | 0.7 | 3e-263 | 1.2 | 9e-16 |
| FOS | 1.2 | 6e-15 | 1.1 | 3e-31 | 0.7 | 1e-32 | 0.7 | 6e-29 | 1.2 | 3e-14 | 1.3 | 4e-161 | 0.9 | 5e-280 | 1.5 | 8e-22 |
| FOSB | 0.5 | 3e-04 | 1.1 | 8e-20 | 1.5 | 6e-69 | 0.6 | 4e-28 | 1.8 | 6e-45 | 1.7 | 4e-191 | 0.4 | 1e-164 | 1.4 | 2e-38 |
| GBP1 |  | NA |  | NA |  | NA |  | NA |  | NA |  | NA |  | NA |  | NA |
| GBP2 | 0.4 | 1 |  | NA |  | NA |  | NA |  | NA |  | NA |  | NA |  | NA |
| IFI30 |  | NA |  | NA |  | NA |  | NA | 0.6 | 1 |  | NA |  | NA |  | NA |
| IFIT1 |  | NA | 0.4 | 1 |  | NA | 0.3 | 1 |  | NA | 0.3 | 0.9 |  | NA |  | NA |
| IFNB1 |  | NA |  | NA |  | NA |  | NA |  | NA |  | NA |  | NA |  | NA |
| IFNG |  | NA |  | NA |  | NA |  | NA |  | NA |  | NA |  | NA |  | NA |
| IL6 |  | NA |  | NA |  | NA |  | NA |  | NA |  | NA |  | NA |  | NA |
| IL6R |  | NA |  | NA |  | NA |  | NA |  | NA |  | NA | 0.3 | 2e-07 |  | NA |
| ISG15 |  | NA | 0.5 | 0.08 |  | NA | 0.3 | 1 | 0.3 | 5e-04 |  | NA |  | NA | 0.3 | 1 |
| JUN |  | NA | 0.7 | 0.01 | 0.6 | 8e-16 | 0.5 | 1e-06 | 0.4 | 1 | 1.0 | 4e-55 | 0.3 | 1e-17 | 0.9 | 2e-05 |
| JUNB |  | NA | 0.5 | 1e-06 | 0.4 | 2e-11 |  | NA |  | NA | 0.3 | 6e-15 | 0.3 | 2e-53 | 0.7 | 7e-07 |
| MX1 |  | NA | 0.4 | 1 | 0.6 | 1e-10 | 0.3 | 1 |  | NA | 0.6 | 5e-06 |  | NA |  | NA |
| MX2 |  | NA |  | NA | 0.3 | 1e-06 |  | NA | -0.3 | 1 |  | NA |  | NA |  | NA |
| PSMB8 |  | NA |  | NA |  | NA |  | NA |  | NA |  | NA |  | NA |  | NA |
| SOCS3 |  | NA |  | NA | 0.6 | 6e-31 | 0.4 | 2e-09 |  | NA | 0.5 | 2e-47 |  | NA | 0.3 | 1 |
| STAT3 |  | NA | 0.3 | 1 | 0.7 | 2e-08 |  | NA |  | NA |  | NA |  | NA |  | NA |
| TYK2 |  | NA |  | NA |  | NA |  | NA |  | NA |  | NA |  | NA |  | NA |

**Supplementary Table 5.** Differential expression of relevant genes related to immune-checkpoint inhibitor myopathy by single-nuclei RNA sequencing. Four biopsies from the cluster ICI-DM were compared with a representative selection of patients with other types of inflammatory myopathy (4 dermatomyositis, 3 antisynthetase syndrome, 6 immune-mediated necrotizing myositis, and 2 inclusion body myositis). Missing values correspond to genes that did not pass the cutoff to be included in the differential expression.

| **Gene** | **T-cells ICI-DM vs. other myositis** | | **FAP cells ICI-DM vs. other myositis** | | **Adipocytes ICI-DM vs. other myositis** | | **Endothelial cells ICI-DM vs. other myositis** | | **Myeloid cells ICI-DM vs. other myositis** | | **Fibroblasts ICI-DM vs. other myositis** | | **Muscle fibers ICI-DM vs. other myositis** | | **Satellite cells ICI-DM vs. other myositis** | |
| --- | --- | --- | --- | --- | --- | --- | --- | --- | --- | --- | --- | --- | --- | --- | --- | --- |
|  | **avg_log2FC** | **p_val_adj** | **avg_log2FC** | **p_val_adj** | **avg_log2FC** | **p_val_adj** | **avg_log2FC** | **p_val_adj** | **avg_log2FC** | **p_val_adj** | **avg_log2FC** | **p_val_adj** | **avg_log2FC** | **p_val_adj** | **avg_log2FC** | **p_val_adj** |
| CEBPB |  | NA |  | NA |  | NA | 0.4 | 2e-17 |  | NA |  | NA | 0.3 | 6e-64 |  | NA |
| EGR1 |  | NA | 1.2 | 9e-29 | 0.6 | 1e-26 | 0.5 | 7e-22 | 0.4 | 5e-16 | 1.1 | 3e-130 | 1.1 | 0 | 1.2 | 1e-21 |
| FOS | 0.7 | 5e-12 | 1.2 | 3e-32 | 0.8 | 1e-32 | 0.7 | 3e-36 | 1.1 | 8e-49 | 1.3 | 3e-200 | 1.3 | 0 | 1.6 | 1e-25 |
| FOSB | 0.4 | 1e-04 | 1.0 | 2e-16 | 1.7 | 3e-78 | 0.3 | 4e-15 | 0.9 | 3e-39 | 1.5 | 4e-180 | 0.3 | 3e-135 | 1.2 | 3e-17 |
| GBP1 |  | NA | 0.6 | 2e-09 | 0.3 | 0.001 | 0.3 | 0.05 | 0.5 | 2e-12 | 0.8 | 9e-42 |  | NA |  | NA |
| GBP2 |  | NA | 0.4 | 1 |  | NA |  | NA | 0.3 | 0.1 |  | NA |  | NA |  | NA |
| IFI30 |  | NA |  | NA |  | NA |  | NA | 0.5 | 0.8 |  | NA |  | NA |  | NA |
| IFIT1 | 1.0 | 2e-25 | 1.1 | 4e-34 | 0.4 | 5e-04 | 1.2 | 9e-32 | 1.0 | 3e-49 | 0.9 | 1e-76 | 1.1 | 0 | 1.4 | 1e-33 |
| IFNB1 |  | NA |  | NA |  | NA |  | NA |  | NA |  | NA |  | NA |  | NA |
| IFNG |  | NA |  | NA |  | NA |  | NA |  | NA |  | NA |  | NA |  | NA |
| IL6 |  | NA |  | NA |  | NA |  | NA |  | NA |  | NA |  | NA |  | NA |
| IL6R |  | NA |  | NA |  | NA |  | NA |  | NA |  | NA |  | NA |  | NA |
| ISG15 | 2.2 | 2e-57 | 2.0 | 6e-53 | 0.8 | 4e-28 | 2.0 | 4e-79 | 2.1 | 5e-119 | 1.8 | 5e-210 | 2.2 | 0 | 2.2 | 2e-47 |
| JUN |  | NA | 1.1 | 7e-15 | 0.5 | 2e-12 | 0.4 | 1e-09 | 0.4 | 1e-04 | 1.1 | 9e-62 | 0.5 | 4e-106 | 0.6 | 0.05 |
| JUNB |  | NA | 0.6 | 2e-08 | 0.4 | 1e-08 |  | NA | 0.3 | 0.06 | 0.5 | 2e-43 | 0.5 | 7e-186 | 0.6 | 2e-06 |
| MX1 | 1.8 | 8e-36 | 1.6 | 2e-43 | 0.5 | 5e-07 | 1.0 | 3e-17 | 2.0 | 1e-73 | 1.4 | 6e-89 | 1.6 | 0 | 1.8 | 4e-40 |
| MX2 | 1.2 | 4e-17 | 1.2 | 1e-20 |  | NA | 1.0 | 7e-19 | 1.2 | 2e-36 | 1.2 | 1e-42 | 0.3 | 4e-106 | 1.2 | 6e-15 |
| PSMB8 |  | NA |  | NA |  | NA |  | NA |  | NA |  | NA |  | NA |  | NA |
| SOCS3 |  | NA | 0.6 | 5e-14 | 0.6 | 1e-29 |  | NA |  | NA | 0.6 | 5e-56 |  | NA | 0.4 | 1 |
| STAT3 |  | NA | 0.7 | 0.04 | 0.6 | 0.5 |  | NA |  | NA | 0.3 | 1 | 0.4 | 8e-39 |  | NA |
| TYK2 |  | NA |  | NA |  | NA |  | NA |  | NA |  | NA |  | NA |  | NA |
